# Transcriptional interference in toehold switch-based RNA circuits

**DOI:** 10.1101/2021.09.29.462367

**Authors:** Elisabeth Falgenhauer, Andrea Mückl, Matthaeus Schwarz-Schilling, Friedrich C. Simmel

**Affiliations:** Physics Department - E14 and ZNN/WSI, TU Munich, Am Coulombwall 4a, 85748 Garching, Germany

**Keywords:** antisense RNA, transcription interference, gene regulation

## Abstract

Gene regulation based on regulatory RNA is an important mechanism in cells and is increasingly used for regulatory circuits in synthetic biology. Toehold switches are rationally designed post-transcriptional riboregulators placed in the 5’ untranslated region of mRNA molecules. In the inactive state of a toehold switch, the ribosome-binding site is inaccessible for the ribosome. In the presence of a trigger RNA molecule protein production is turned on. Using antisense RNA against trigger molecules (anti-trigger RNA), gene expression can also be switched off again. We here study the utility and regulatory effect of antisense transcription in this context, which enables a particularly compact circuit design. Our circuits utilize two inducible promoters that separately regulate trigger and anti-trigger transcription, whereas their cognate toehold switch, regulating expression of a reporter protein, is transcribed from a constitutive promoter. We explore various design options for the arrangement of the promoters and demonstrate that the resulting dynamic behavior is strongly influenced by transcriptional interference (TI) effects, leading to more than four-fold differences in expression levels. Our experimental results are consistent with previous findings that enhanced local RNA polymerase concentrations due to active promoters in close proximity lead to an increase in transcriptional activity of the strongest promoter in the circuits. Based on this insight, we selected optimum promoter designs and arrangements for the realization of a genetic circuit comprised of two toehold switches, two triggers and two anti-triggers that function as a post-transcriptional RNA regulatory exclusive OR (XOR) gate.

## INTRODUCTION

RNA-based regulatory mechanisms are an important component of biological gene regulation networks. For instance, small RNA molecules (sRNAs) modulate gene expression levels in bacteria by binding to complementary sequences in mRNA molecules and thus influence their lifetime. Similarly, microRNAs are involved in fine-tuning expression levels through RNA interference in eukaryotes. Other regulatory roles for RNA are found in sensing of metabolites via riboswitches ^1^, bacterial defense mechanisms (i.e., CRISPR) ^2, 3^ or the regulation of quorum sensing ^4^.

Regulatory RNAs are increasingly used also as components for synthetic biological circuits ^5^. As their function is mainly determined by predictable nucleic acid base-pairing interactions, they can be engineered more straightforwardly than protein-based regulators. RNA regulation can be exerted at various stages during gene expression – during transcription (e.g., by modulation of transcriptional termination/anti-termination), post-transcriptionally (e.g., by controlling translation initiation), or post-translationally (e.g., by controlling the degradation of mRNA) ^6, 7^.

One of the first examples for a synthetic post-transcriptional riboregulator was developed by Isaacs et al. ^8^. In their riboregulator concept, the ribosome binding site (RBS) present in the 5’ untranslated region (5’ UTR) of an mRNA molecule was sequestered within the stem of a stable RNA hairpin structure – similarly as found in riboswitches -, which resulted in a strong reduction of translation initiation and thus protein expression. In the presence of a trans-activating RNA molecule that was partly complementary to the RNA hairpin sequence, the stem of the hairpin was opened and protein translation activated.

Building upon this work, Green and co-workers ^9^ later developed a rationally designed riboregulator with considerably improved switching characteristics. In their design, the RBS is sequestered in the loop of an RNA hairpin, while the start codon is included as a bulge in its stem (cf. Figure 1A). Notably, strand invasion by the trans-activating trigger RNA is facilitated by a single-stranded toehold preceding the hairpin-stem loop structure. The best-performing toehold switch was found to have a remarkable ON/OFF ratio of ≈ 600, two orders of magnitude above previous RNA-based gene activation processes. Toehold switches can be engineered to recognize different trigger RNAs as inputs, which already has led to several applications in medical diagnostics ^10^.

**Figure 1:**
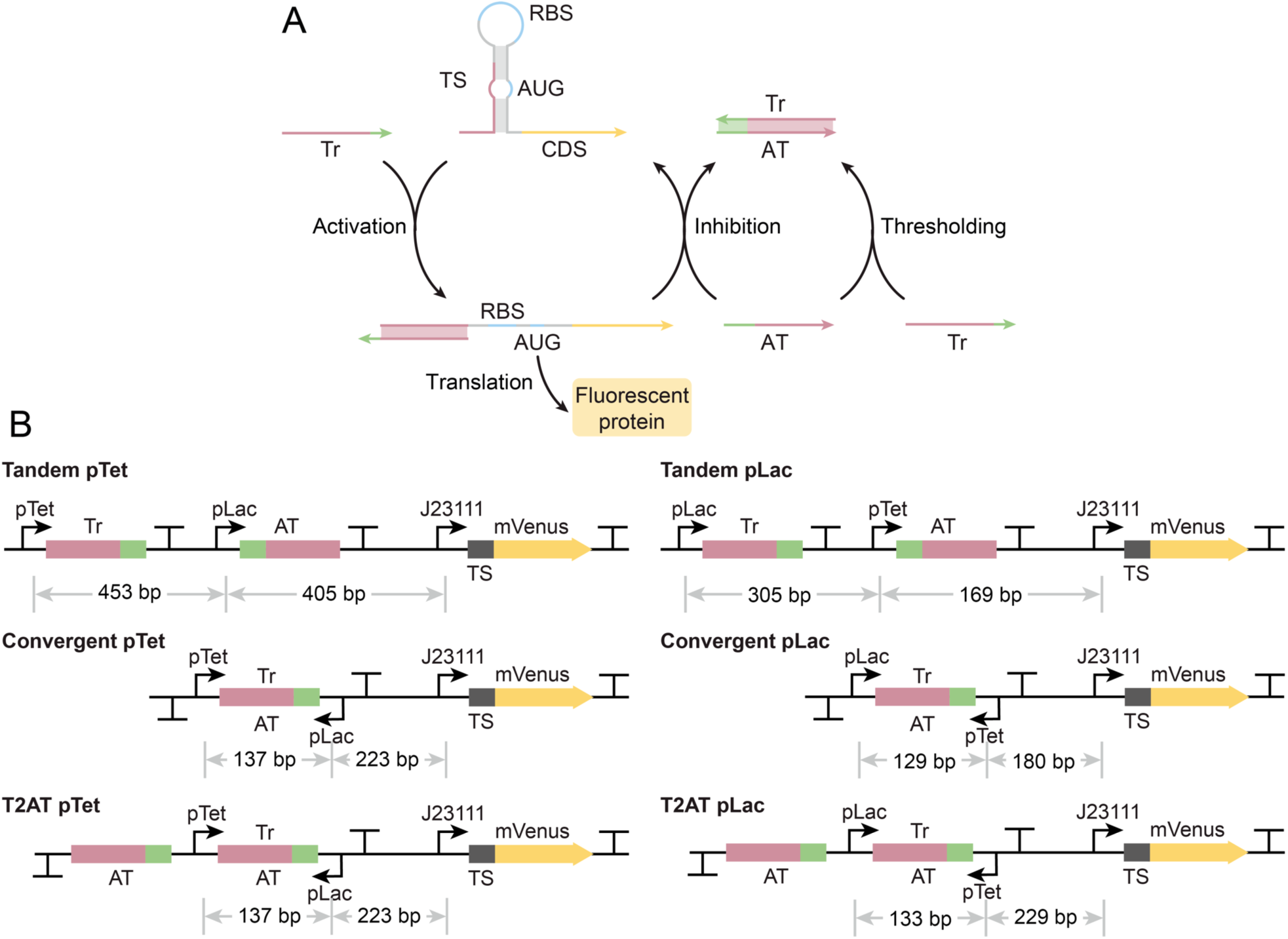
(A) In the inactive state of a TS, the ribosome-binding site is hidden in an RNA hairpin loop and is thus inaccessible for the ribosome. Tr binds the toehold, opens the secondary structure via a strand displacement process and therefore activates gene expression. AT can counter-act Tr by sequestering excess Tr via direct hybridization (thresholding), or by removing Tr from activated trigger-toehold switch complex via toehold-mediated strand displacement. (B) Promoter arrangements of the pTet designs (left). A tandem design without overlapping transcription cassettes was tested as reference system. In the convergent design the trigger/anti-trigger sequence is embedded between the two promoters, the transcription terminators are downstream of the converging promoters. A second convergent promoter design was cloned, which is extended in one direction by a ribozyme and a second anti-trigger sequence. In contrast to the convergent pTet and T2AT pTet design, the distances between adjacent promoters are large in the tandem pTet design. (right) All promoter arrangements were also tested with exchanged promoters referred to as pLac designs or tandem pLac, convergent pLac and T2AT pLac. The distance between pLac and pTet is large in the tandem pLac design, the distances between all other adjacent promoters are small.

In an alternative approach, Lucks and coworkers engineered naturally occurring transcriptional attenuators to obtain a set of inducible riboregulators termed STAR (small transcription activating RNA) regulators. In STAR mechanisms, either a terminator or an anti-anti-terminator sequence is put upstream of a gene sequence of interest. Inducible expression of an antisense (STAR) molecule then induces a conformational change in the terminator (or anti-anti-terminator), resulting in anti-termination and thus activation of transcription ^11, 12^. A second level of antisense RNA molecules (anti-STAR RNA) even allows the regulation of multiple genes in a reversible and orthogonal manner ^13^. Antisense RNA was later also applied in the context of toehold switch-based logic gates ^14^, where it was used to implement logical negation (NOT gates). Besides the sequence of the regulatory RNA molecules themselves, there are other important design aspects in genetic circuits such as the strengths of the promoters and their arrangement in the genome. In nature, small antisense RNA molecules are encoded either as *cis-* or *trans*-acting regulatory sequences ^7, 15^. For example, sense and antisense RNA may be produced from the same stretch of DNA (in *cis*) via transcription from two distinct promoters flanking the sequence, with transcription directions pointing face-to-face (“convergent promoters”), from overlapping promoters facing in opposite directions (“divergent promoters”), or by promoter arrangements with co-directional transcription (“tandem promoters”). Sneppen and coworkers^16^ found 54 non-overlapping convergent promoters with a distance between 40-200 bp in *E. coli*, which comprise only a small fraction compared to other tandem or divergent promoter arrangements. Convergent promoter arrangements embracing larger sequence stretches are even less frequent. However, convergent designs not only allow a compact genetic design but also allow processitivity (uninterrupted transcription) control via transcriptional interference mechanisms ^17, 18^.

Transcriptional interference (TI) is an effect resulting from the influence of strong (“aggressive”) on weak (“sensitive”) promoters in cis^19, 20^. Transcriptional interference is surmised to fulfill important functions in many cellular processes, both in prokaryotes and eukaryotes. This involves, e.g., initiation of plasmid replication^19^, or regulatory roles during embryonic development in *Drosophila*^21^. TI may also play an evolutionary role through its effect in head-on collisions between replisomes and RNAPs, which potentially increases the evolvability of convergently oriented genes through mutagenesis^22^. Moreover, several metabolic engineering tasks have been successfully implemented via convergent promoter designs^23-25^. Several TI mechanisms are known. The passage of an RNA polymerase (RNAP) over a promoter can either disrupt the initiation process (sitting duck interference) or occlude the RNAPs from binding to the promoter site (occlusion) depending on whether the promoter has a slow RNAP escape rate or a small RNAP binding rate, respectively. Head-on collisions of elongating RNAPs can cause stalling of the RNAPs or a premature termination of one or both polymerases. This effect is more likely for larger promoter distances and promoter activity. A cooperative push-through of a leading RNAP with several trailing RNAPs is possible^20, 26-28^. A similar situation arises for DNA binding proteins like LacI or TetR, which act as a roadblock for an elongating RNAP. Here it was shown that a polymerase blocked by a lac repressor protein (LacI) can be removed by the transcription-repair coupling factor Mfd from the DNA^29^. It is also possible that a leading polymerase followed by trailing RNAPs can cooperatively read through a roadblock. However, such a cooperative dislodgment occurs only when a high RNAP flux is provided by a strong promoter.

Also the relative distance of promoters to each other has been shown to have an effect. Active promoters increase the local RNAP concentration around their promoter sites. For small promoter distances, this can result in a simultaneous upregulation of a strong promoter and downregulation of a weak promoter in close proximity as the strong promoter benefits from the increased local RNAP concentration caused by both active promoters and is able to recruit the major portion of the RNAPs^30^.

It is conceivable that some of these features may also prove useful in the context of synthetic gene circuits. Recently, transcriptional interference has gained the interest of synthetic biologists as a potentially engineerable tool to manipulate RNAP traffic on DNA^30^ and to tune gene expression levels^31^. Using a variety of synthetic gene constructs, Bordoy *et al*. investigated TI effects of convergent promoter designs in detail to assess their tunability and used them to regulate gene expression over several order of magnitude. On the downside, unwanted transcriptional interference also poses a design challenge for synthetic circuits of increasing complexity^32^.

In the following, we focus on genetic constructs with toehold switches *cis* encoded with their triggers and anti-triggers. In detail we activate the expression of a fluorescent protein from a toehold switch by inducing the production of trigger RNA from an inducible promoter. Counteracting this activation process, the production of an “anti-trigger” RNA with a sequence complementary to the trigger is used to buffer away trigger molecules and correspondingly reduce protein production. We explore the influence of promoter orientation on the performance of our gene circuits in the context of transcriptional interference. Based on our results, we combine the best performing designs, and use them to implement a toehold switch-based XOR logic gate in *E*.*coli*.

## RESULTS AND DISCUSSION

### Modulation of toehold switching via antisense trigger molecules

The fundamental circuit motif utilized in our work comprises a toehold switch together with its cognate trigger and anti-trigger RNA molecules. As shown in Figure 1A, Tr can disrupt the stem of the TS via toehold-mediated strand invasion, which exposes the RBS and start codon initially sequestered in the hairpin and thus activates translation of the mRNA sequence (which, in this case, codes for a fluorescent protein). AT can counteract the Tr (and thus reduce reporter expression) by sequestering excess Tr via direct hybridization, or by removing Tr from activated trigger-toehold switch complex via toehold-mediated strand displacement.

As an initial characterization of this circuit motif, we purified TS RNA regulating translation of a fluorescent protein (CFP), the corresponding Tr RNA and anti-trigger RNA and tested them in a cell-free gene expression system (Figure S1 A and B). For a fixed TS concentration of 90 nM, we observed a proportional increase in the fluorescence end levels of the CFP reporter protein for increasing trigger concentrations up to 90 nM. For Tr concentrations over 200 nM we observed a saturation in the signal end levels. In an anti-trigger screening experiment, we kept the TS and Tr concentration constant at 90 nM and 200 nM, respectively. Increasing anti-trigger concentrations decreased the fluorescence end levels steadily also for AT concentrations above 200 nM. An extrapolation of the results shown in Fig. S1 suggest that around 400 nM of anti-trigger would be necessary to reduce the output signal to zero. We further confirmed the functionality of TS, Tr and AT in an electrophoretic mobility shift assay (Figure S1C) as hybridization products can be observed for Tr and TS and also for Tr and AT. Undesired hybridization between TS and AT was not observed.

### Promoter arrangements for trigger and anti-trigger RNA

We next tested the function of triggers and anti-triggers on the toehold switch in *E*.*coli* by cloning the corresponding genetic elements onto a single plasmid (“in cis”). Similar as in naturally occurring cis-encoded antisense regulation, we encoded the sequence complementary Tr and AT molecules using two convergent promoters flanking the trigger sequence. As alternative designs we chose tandem promoters where we terminated the first transcript upstream of the second promoter to prevent TI effects based on collisions or passage of the RNAPs over the promoter region. As our cell-free characterization experiments indicated a beneficial effect of excessive AT concentrations, we also cloned a convergent promoter design (T2AT), in which antisense transcription would result in the generation of two anti-trigger molecules, while sense transcription would only produce a single trigger. The two anti-triggers on the antisense transcript were separated from each other post-transcriptionally using a self-cleaving ribozyme as an insulator^33^. The functionality of this 2AT design was also confirmed in cell-free expression experiments using the PURExpress system (Figure S2).

While the toehold switch controlling the expression of a YFP reporter protein was transcribed constitutively (iGEM part J23111, Figure 1B), we regulated the transcription of trigger RNA using an aTc inducible pTet promoter (pLTet-O1, referred to as pTet) and the transcription of anti-trigger using an IPTG inducible pLLac-O1 promoter (referred to as pLac). We also tested the same promoter arrangements as in the “pTet-Tr designs” with exchanged promoters (referred to as pLac-Tr designs). We expected a difference in reporter expression due to the differences in promoter strength and the resulting TI effects. The promoter strength of J23111 (Figure S3) is about 300% higher compared to pTet (Figure S3) and pTet is about 30% stronger than pLac (Figure S4).

The corresponding circuit plasmids were each transformed into *E*.*coli* strain DH5a Z1, which endogenously codes for LacI and TetR, and were then induced with 42 combinations of IPTG vs. aTc concentrations. In the following we use a shorthand notation for the inducer combinations, where Ixay denotes concentrations of x mM IPTG and y ng/ml aTc. The fluorescence intensity normalized by the absorbance signal (FI/Abs) after 540 min of induction was compared for fixed AT inducer concentrations and further analyzed by fitting the Hill curve

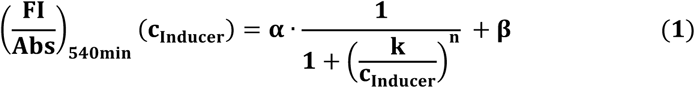

to the data, where α denotes the scaling factor (fold change), β the offset of the curve (baseline expression), k the induction threshold and n the Hill coefficient (see Methods section for details).

### Comparison of the tandem reference systems

In the tandem pTet design Tr is transcribed from a Tet promoter, AT from a Lac promoter, while the toehold switch controls the expression of an mVenus reporter and is transcribed from a J23111 promoter. All promoters point in the same direction and are separated by at least 400 bp (Figure 2A), which is expected to reduce local RNAP concentration effects to a minimum. In addition, the transcripts are terminated upstream of the following promoters with a strong terminator (iGEM part B0015, termination strength >98%) or a combination of terminators B1002 (termination strength 98%) and B1006 (termination strength 99%), in order to avoid collision events between elongating RNAPs and transcriptional read through. The maximum signal for the I0a200 inducer combination is around 6400 a.u., and the signal can be repressed about 3.3-fold via anti-trigger expression (Figure 2B, OFF state defined as I0.5a200).

**Figure 2:**
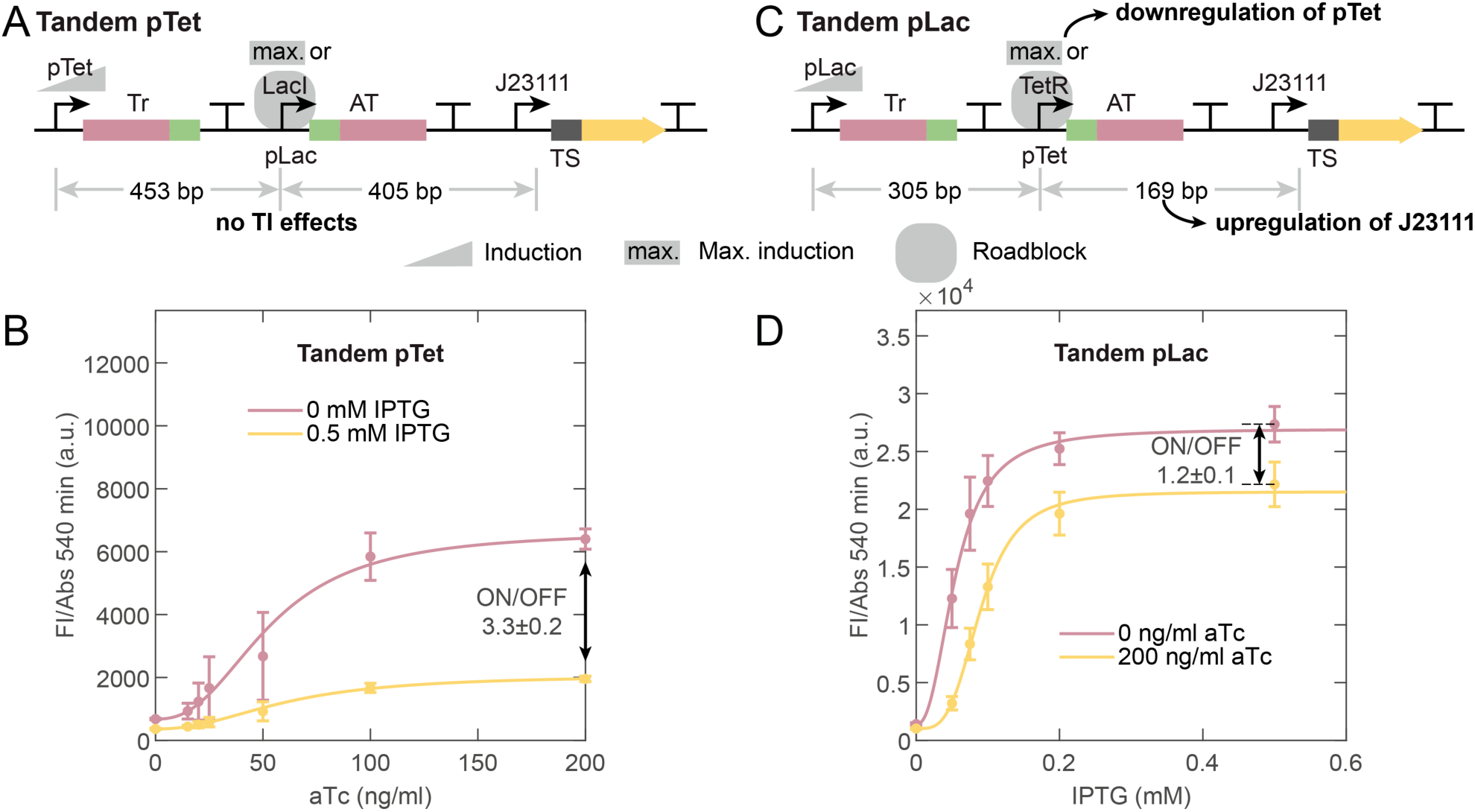
Comparison of the reference systems. (A) All promoter distances in the tandem pTet design are large. We compare induction of pTet at a fully induced (0.5 mM IPTG) or uninduced (0 mM IPTG) pLac promoter. (B) Results for the experiment described in (A). The maximum signal is around 6400 a.u. and the ON/OFF ratio is around 3.3. (C) The promoter distance between pTet und J23111 is small in the tandem pLac design. We compare induction of pLac at a fully induced (200 ng/ml aTc) or uninduced (0 ng/ml aTc) pTet promoter. (D) Results for the experiment described in (B). The maximum signal is more than 4 times higher compared to the tandem pTet design, the ON/OFF ratio is about 2.8 times smaller.

In the tandem pLac design the distance between the inducible promoters is 305 bp, which is also assumed to be sufficiently ‘far’, but pTet and pLac are exchanged. In contrast to the tandem pTet design we deleted B1006 to reduce the distance between pLac and J23111 to 169 bp (Figure 2C). The exchange of the promoters with 30% difference in promoter strength should cause a reduced maximum signal at I0.5a0 and a higher ON/OFF ratio when comparing I0.5a0/I0.5a200. However, the small distance between pTet and J23111 appears to cause a local RNAP concentration effect, which upregulates J23111 and simultaneously downregulates pTet. For 0 ng/ml aTc it is plausible, that RNAPs released after Tr transcription might be also utilized by the strong neighboring J23111 promoter, resulting again in enhanced production of toehold switches^34^. The result is a 4.3-fold increase in the maximum signal compared to tandem pTet, but also a decrease in the ON/OFF ratio to merely about 1.2 (Figure 2D).

For a better comparability of all other (convergent) circuit designs we kept the promoter distances between J23111 and the closest upstream promoter ‘short’, i.e., at a distance between 180 and 229 bp (the distances are measured between the centers of the promoter sequences to be independent of promoter orientation). Also the distances between the convergent promoters was kept short at a distance of 129-137 bp.

### Induction curves in absence of anti-trigger induction

We next investigated the TI effects on the trigger induction curves in the absence of an inducer for anti-trigger transcription. Repressor proteins bound to a converging promoter can cause potential roadblock effects. This effect is not expected for the tandem designs as the first transcript is terminated upstream of the second promoter. Previous studies have shown that roadblocks can either cause premature transcriptional termination or can be cooperatively dislodged by several polymerases, if the RNAP flux is high enough (high promoter strength). TetR has been found to cause a stronger roadblock than LacI. In contrast to a study by Bordoy et al. ^30^, whose transcripts overlapped just for a part of their sequence, for our designs (Fig. 1B) we would expect to obtain fully functional trigger RNA sequences even in the case of transcriptional termination at the roadblock as the full trigger sequence is upstream of the roadblock.

In the pTet-Tr designs, in the absence of IPTG a LacI roadblock interferes with trigger transcription starting at the Tet promoter. For maximum induction, the convergent pTet design reaches a signal which is about 58% higher than that for the tandem pTet design, which is terminated upstream of the roadblock. Also the T2AT pTet design is found to be upregulated by 16% (Figure 3B). It is conceivable, that the premature termination at the roadblock increases the effective RNAP concentration close to the promoter as the RNAP becomes more readily available for another round of transcription, resulting in an upregulation of the Tet promoter in the convergent designs.

**Figure 3:**
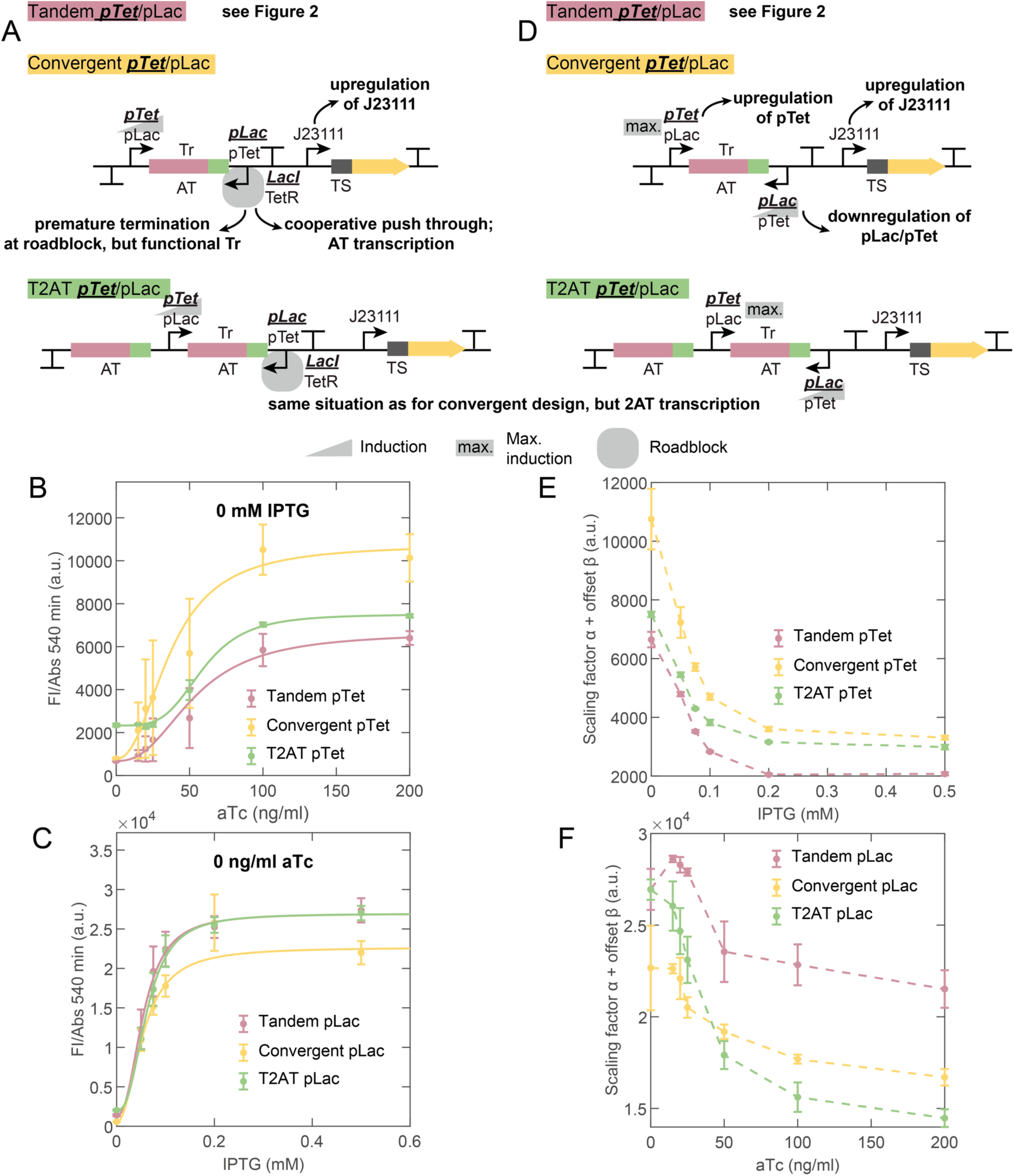
(A) The induction curves for the Tet promoter in the presence of the LacI roadblock are compared for all pTet designs. Analogously, the induction curve for the Lac promoter in the presence of the TetR roadblock is compared. Just for the tandem designs the roadblock is downstream of a transcription terminator. The potential TI effects are annotated. (B) Induction curve at 0 mM IPTG for the pTet-Tr designs. The convergent pTet and T2AT pTet design are upregulated compared to the tandem pTet design. (C) Induction curve at 0 ng/ml aTc for the pLac-Tr designs. All designs show almost the same induction curves. (D) Repression capability of the designs is tested by induction of AT transcription at maximum trigger transcription. The potential TI effects are annotated. (E) The corrected scaling factor (scaling factor α + offset β) for the pTet-Tr designs is a measure of the repression capability of the designs. The convergent pTet design shows the highest, tandem pTet the lowest α+β values. (F) Corrected scaling factor for the pLac-Tr designs. tandem pLac shows a small increase followed by a decrease in α+β, the convergent and T2AT design show a stronger decrease.

Analogously, we focus on the IPTG induction curves for the pLac-Tr designs in the absence of aTc, which involves a TetR roadblock (Figure 3C). The induction curves for tandem pLac, convergent pLac and T2AT pLac almost overlap over the full IPTG concentration range, but as a result of the higher variance of the I0.2a0 and I0.5a0 samples, the convergent pLac Hill fit is shifted to smaller values. The absence of any upregulation is surprising as TetR is expected to cause an even stronger roadblock than LacI^30^.

A closer look at the promoter distances between J23111 and the upstream promoter suggests an alternative mechanism: The upregulation of the convergent pTet and T2AT pTet designs in comparison to tandem pTet is likely not caused by the roadblock and a corresponding upregulation of the Tet promoter (and thus higher Trigger availability), but probably is a consequence of the shorter 223 bp promoter distance between pLac and J23111 compared to the 405 bp in the tandem pTet design. Thus RNAPs released at the roadblock might be utilized by the strong neighboring J23111 promoter^34^, resulting in enhanced production of toehold switches.

The observations for the IPTG induction curves for the pLac-design are in line with this interpretation, as the absence of upregulation is consistent with the constant promoter distance between pTet and J23111 for all pLac-designs. The upregulation effect on J23111 of a close pLac promoter is much smaller than that of a close pTet promoter, as the maximum signal obtained for the convergent pTet design is still by a factor of 2.7 smaller compared to the tandem pLac design.

The shift of T2AT pTet to smaller values can be explained as the result of cooperative unbinding of the roadblock by transcribing polymerases, as described above, and the subsequent transcription of two anti-triggers (instead of one for the convergent pTet design). This effect was not observed for the pLac designs, as the TetR roadblock is stronger and the promoter strength of pLac might be too small to accumulate enough RNAP for a cooperative push-through.

### Influence on the repressive effect of anti-trigger transcription

To analyze the repression capability of the designs, we focus on the ON/OFF ratios defined as the maximum induced state divided by the state with both inducers at their maximum tested concentration. As this ratio merely conveys information about the relative position of two points, we also focus on the variation of the sum of the scaling factor α and the offset β of the Hill fits (see Equation 1), referred to as corrected scaling factor. This parameter shows the effect of increasing amounts of AT (induction of AT promoter) at a maximum trigger concentration (fully induced promoter). The parameter α+β decreases for increasing anti-trigger inducer concentrations for both tandem designs because of the expected thresholding and inhibition effect of transcribed AT.

We observe higher corrected scaling factor values for the convergent pTet design compared to the tandem pTet design (Figure 3E). The Lac promoter is inhibited in AT transcription by the Tet promoter by a combination of collisions and passage effects. In addition, pLac is inhibited from both sides by local RNAP concentration effects. The activity of both adjacent promoters can be upregulated causing a higher TS and Tr availability and a constant ON/OFF ratio (Table S1). Corrected scaling factor values of T2AT pTet are in between the other two designs. The double AT concentration can partly counteract the inhibition on the Lac promoter. However, the ON/OFF ratio is about 20% smaller for this design.

For the tandem pLac design, we observe a small initial increase of the corrected scaling factor in response to elevated aTc concentrations, which is followed by a subsequent decrease for higher concentrations. For the convergent pLac design we observe an initially constant corrected scaling factor and a subsequent decrease, while for the T2AT pLac design we observe the steepest decrease in the α+β value (Figure 3F). The ON/OFF ratios are in a range between 1.2 (tandem) and 1.9 (T2AT) and therefore smaller compared to the pTet-Tr designs. The close distance of J23111 repressed pTet activity significantly, so the interaction between these two promoters is in both directions (inhibition of pTet and upregulation of J23111) higher than for pLac and J23111. The induction of pTet close to J23111 regulates J23111 further up, but the increasing AT transcription diminishes this effect quickly. For the convergent design this effect is less pronounced as the activated pTet also downregulates Tr transcription starting at pLac, again by a combination of collision, passage and local RNAP concentration effects. The T2AT design profits from a combination of TI effects and expression of a double amount of AT, and therefore has a 58% higher ON/OFF ratio compared to the tandem design.

### Effects on Hill coefficient and dissociation constant

We also study the dependence of the Hill coefficients n and dissociation constants k of our Hill fits (see Equation 1) on AT induction. These two parameters provide information about the change in sensitivity to trigger induction (slope of the trigger induction curve and point of half maximum induction). Bordoy et al. found that the TI effect mainly influences the Hill scaling factor, while n and k stayed almost constant. In contrast, in our experiments we observe an increase in k for increasing AT inducer concentrations for all designs except tandem pTet in the range of 31% - 64% (Figure S6 and S8). With only 14%, the induction threshold of tandem pTet shows the lowest increase of all designs. The upregulation of J23111 in all designs except tandem pTet results in a shift of the ratio between TS and Tr to the benefit of TS, which makes the system more sensitive to AT and causes an increase of k. As this upregulation on J23111 and downregulation on pTet is not present in the tandem pTet design, we observe the smallest increase.

By contrast, the Hill coefficients stay roughly constant over the AT inducer screening range for all designs. The highest Hill coefficients were observed for T2AT pTet (almost 4) and tandem pLac (around 3). All other designs show a Hill coefficient around n=2.5. For T2AT pTet this is mainly caused by the higher offset for the I0a0 condition, whereas for tandem pLac it is a result of the high degree of upregulation as discussed before.

In summary, our results on the induction curves and the repression capability suggest that the dominant transcriptional interference effect in our circuit designs is caused by local RNAP concentration effects due to the small distance of J23111 to our inducible promoters. We also observed effects that could be explained by a cooperative push of RNAPs through a roadblock, and a combined collision and passage effect. We were not able to detect effects, which could be caused by premature termination of transcripts based on roadblock effects, which was an expected result as our convergent designs always enclosed the full trigger sequence. The pLac designs benefitted most from the TI effects and showed up to four-fold higher maximum signals (induction curves in absence of AT) compared to the pTet designs even though Tr transcription was regulated by the weaker pLac promoter. The highest ON/OFF ratio among all pLac designs was obtained for T2AT pLac, which benefited from the double AT transcript (≈ 1.9). The highest maximum signal in the pTet-Tr designs in the absence of anti-trigger was observed for the convergent pTet design, which benefited from an upregulation effect on J23111. This design, together with tandem pTet, displayed the best ON/OFF ratio in our study (≈ 3.2).

### Implementation of an XOR gate based on the optimum anti-trigger design

Based on these results, we designed a logic gene circuit with the function of an exclusive OR (XOR) gate, for which we utilized two separate toehold switches together with their triggers and anti-triggers (Figure 4A and B). One trigger/anti-trigger pair was based on our T2AT pLac design, while the other utilized a convergent pTet design. The first toehold switch (XOR YFP) was identical to that used to investigate the TI effects above, while the second switch (XOR CFP) was designed with an altered sequence (version 2 ^9^), and the reporter protein was changed to CFP. As a result we obtained a gene circuit, in which Tr1 and AT2 are expressed under IPTG induction, while Tr2 and AT1 are expressed by aTc induction, i.e., each inducer activates one of the toehold switches and simultaneously represses the other.

**Figure 4:**
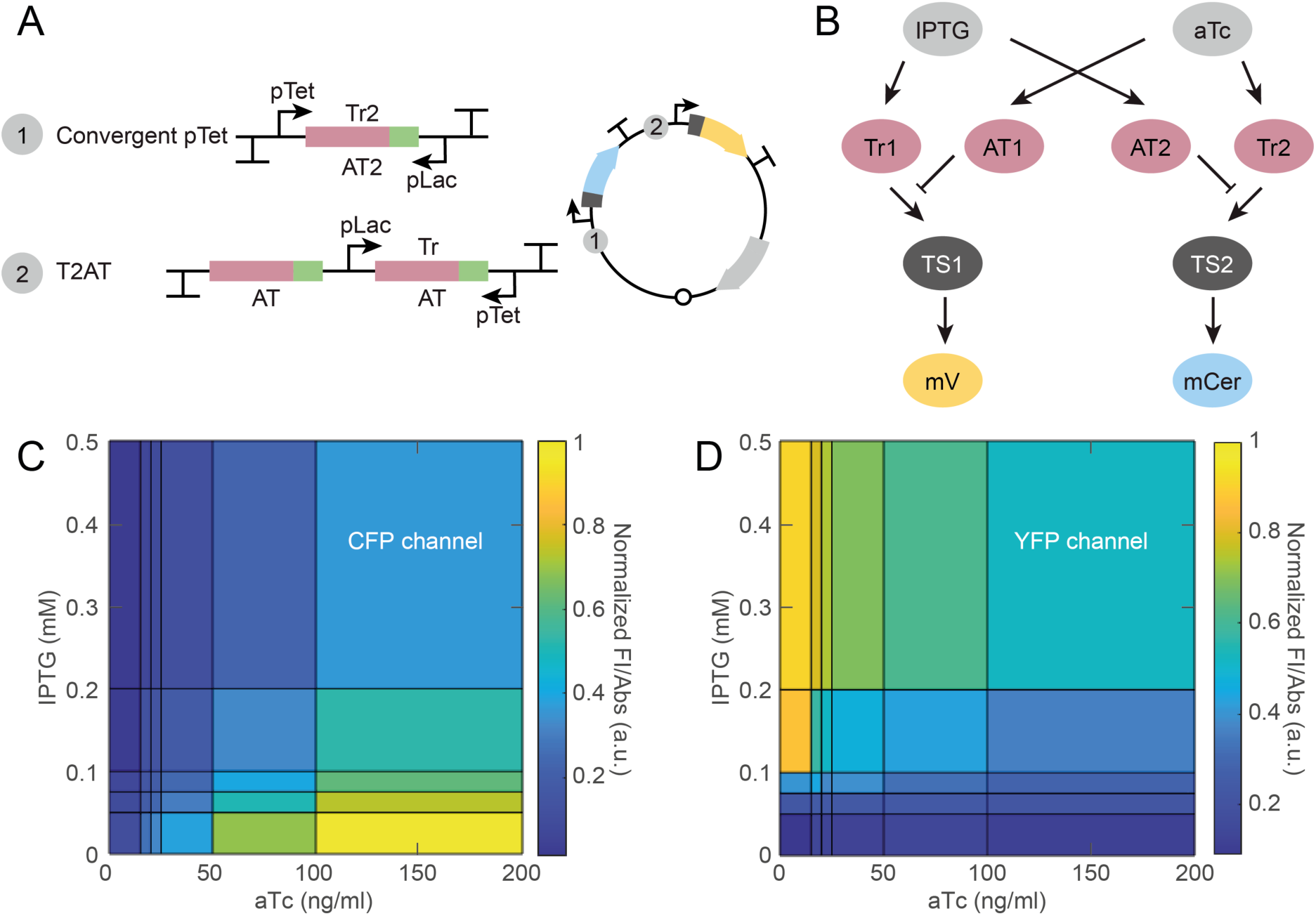
XOR gate. (A) A convergent pTet design based on TS2, Tr2, AT2 and a CFP reporter is combined with a T2AT pLac designs to get a XOR logic gate. (B) Regulation schematics. Tr1 and AT2 are transcribed under IPTG induction, Tr2 and AT1 under aTc induction and regulate TS1 and TS2. A YFP is expressed if TS1 and a CFP is expressed if TS2 is active. (C) Heat map for the YFP reporter of the XOR gate after 540 min of induction. A region around high IPTG but low aTc concentrations shows a high expression level of the YFP reporter. (D) Heat map for the CFP reporter of the XOR gate after 540 min of induction. For high aTc but low IPTG concentrations the CFP reporter is expressed to a high level.

The gene regulation function of the XOR gate (i.e., reporter expression as a function of the two input inducers) has two regions, where one of the reporters is dominant, while the other is expressed at a very low level, and a third region, where both reporters are co-expressed at similar, but relatively low levels (Figure 4C and D). Even though the overall signal levels are smaller (most likely due to the higher metabolic load of the more complex circuit on the bacteria), the two linked toehold switch systems display almost the same ON/OFF ratios (1.7±0.1 for XOR YFP and 3.2±0.2 for XOR CFP) as when studied in isolation. Due to the different reporter proteins, we normalized the fluorescence signals to the maximum at I0.5a0 and I0a200, respectively. In order to demonstrate dynamic switching between the different logic states of the XOR gate, we chose four distinct points of the gene regulation landscape (corresponding to the logical input combinations 00, 01, 10, 11) as the starting or end points of a switching process and characterized all 12 possible transition pathways between them using plate reader measurements (Figure 5 and Figures S11-13). For example, Figure 5A and B compare the reporter signals for the switching processes from I0a0 to I0.5a0 and back. As expected, the YFP signal increases and the CFP signal stays at a low background level. Back switching causes a signal decrease in the YFP channel and a small increase in the CFP signal. As our reporter proteins were not equipped with a degradation tag, the signal decrease is caused by dilution due to bacterial growth. In comparison to the OFF signal at 0 min in Figure 5A, the end levels of both channels are higher, but can be well distinguished from the ON states and also from the intermediate state at I0.5a200. Figure 5C shows switching between the dominant regions. After 120 min the YFP reporter signal exceeds the CFP signal, which again decreases due to bacterial growth. The addition of 0.5 mM IPTG to a culture, which was already induced with 200 ng/ml aTc, causes a transition of the system to a state with co-expressed reporters at an intermediate level, which is reached within less than 250 min (Figure 5D).

**Figure 5:**
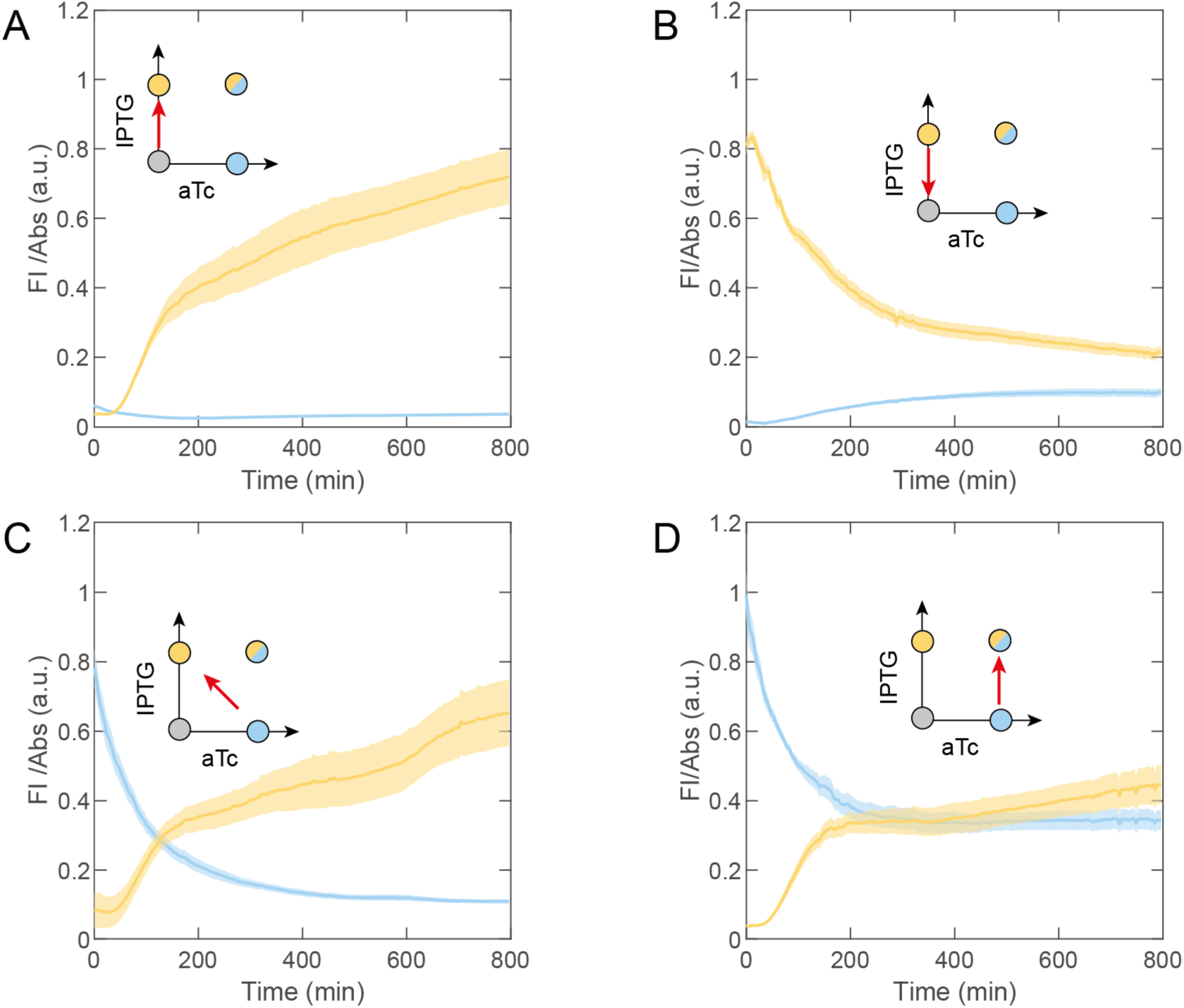
XOR switching experiments. (A) Switching from I0a0 to I0.5a0 causes a signal increase in the YFP channel but not in the CFP channel. (B) The system can be switched off again. (C) Transition between the dominant regions (I0a200 to I0.5a0). (D) Transition to the intermediate state at I0.5a200 starting at I0a200.

## CONCLUSION

We used toehold switches in combination with activating trigger and deactivating anti-trigger RNAs to study transcriptional interference effects for different arrangements of inducible promoters. We observed mainly local RNAP concentration effects due to a small distance between the strong constitutive J23111 promoter controlling transcription of the toehold switch and our inducible promoters. We also observed effects that can be explained by a cooperative push-through of multiple RNAPs through a roadblock. We avoided the production of unfunctional Tr and AT by premature termination at roadblocks by putting their full functional sequence upstream of the roadblock, which was possible due to the sequence complementarity of Tr and AT and the short transcript length. The pLac designs benefitted most from the TI effects and showed up to four-fold higher maximum signals compared to the pTet designs, even though trigger transcription was regulated by the weaker pLac promoter. The best designs were assembled to a XOR logic gate, which shows fast switching kinetics.

The use of RNA trigger molecules and their exactly sequence-complementary anti-triggers within toehold switch circuits naturally suggests their transcription from the same DNA template using convergent promoters. This leads to particularly compact circuit designs, but comes with the price of transcriptional interference effects. Depending on the extent of such effects, widely varying gene expression responses can be generated. As demonstrated in our work, a solid understanding of these TI effects supports their rational implementation within toehold switch circuits and can strongly enhance their performance.

## METHODS

### Plasmids

Plasmids were cloned using a restriction-ligation protocol. Linear gene fragments were ordered from IDT and cloned into the target vectors (pSB1A3, pSB3C5). Final plasmid sequences are listed in Table S2-8.

### Bacterial strains and culture conditions

Plasmids were transformed to DH5alpha Z1. Cells from glycerol stocks were grown in 5 ml Luria-Bertani (LB) medium containing selective antibiotics and 20 mM glucose at 37°C overnight and 250 rpm. The following day, cells were diluted 1:100 in M9 medium for plate reader experiments and were grown to an OD of 0.3, keeping the antibiotic conditions the same.

### Plate reader experiments

Cells were transferred to 384 well plates (ibidi) and induced with IPTG (0mM, 50 nM, 75 nM, 0.1 mM, 0.2 mM and 0.5 mM) and aTc (0 ng/ml, 15 ng/ml, 20 ng/ml, 25 ng/ml, 50 ng/ml, 100 ng/ml and 200 ng/ml). Fluorescence and absorbance time traces were recorded using a plate reader (Fluostar Omega – BMG Labtech). Blank values were subtracted, the fluorescence levels at 540 min were normalized to the corresponding absorbance values (FI/Abs) and plotted against the inducer concentration of the promoter regulating Tr expression for fix AT inducer concentrations (Figure S5 and S7). The error bars show the standard error of 3 biological replicates Hill curves were fitted to the data and the course of the fitting parameters a (scaling factor), b (offset), K (dissociation constant) and n (Hill coefficient) were plotted against the anti-trigger inducer concentration (Figure S6 and S8).

### XOR switching experiments

DH5alpha Z1 cells containing the XOR plasmid were cultured overnight in four separate culture tubes with LB + glucose medium and the selective antibiotic (Cm). The cultures were induced with either of four inducer combinations of 0 mM or 1 mM IPTG against 0 ng/ml or 0.1 ng/ml aTc. Cells were diluted 1:100 in M9 medium keeping the inducer conditions the same. At OD 0.3 the cultures were centrifuged (3000 g, 3 min, 4°C) and resuspended in 1/10 of the starting volume. Inducer conditions were kept constant or changed to one of the three other combinations by diluting 30 μl of bacteria in 270 μl M9 medium containing the corresponding inducers. Fluorescence and absorbance time traces were recorded using a plate reader. The FI/Abs time traces are shown in Figure 5 and S11-12, and just the start and end levels are shown in Figure S13 for better comparison.

### Cell-free expression tests

Transfer functions of Tr and AT were studied using purified RNA. Separate in-vitro transcripts were purified using a standard phenol-chloroform precipitation. RNA concentration was estimated based on a denaturing UREA-PAGE. A cell-free gene expression test was performed as described previously ^35^.

Functionality of the modified anti-trigger (2AT = AT-Ribozyme-AT) was proven in a cell-free test using a linear DNA fragment coding for a T7 promoter and the desired RNA sequence (TS1-YFP, trigger or modified and unmodified anti-trigger). The reaction was set up using a commercial kit (PURExpress NEB) according to user guidelines. Fluorescence time traces were recorded using a plate reader (Fluostar Omega – BMG Labtec).

## ASSOCIATED CONTENT

**Supporting Information**

## AUTHOR INFORMATION

### Notes

The authors declare no competing financial interest.

## ACKNOWLEDGMENTS

We gratefully acknowledge financial support by the BMBF through the ERASYNBIO network (project UNACS, grant no. 031L0011) and the European Research Council (grant agreement no., project AEDNA). M. Schwarz-Schilling is supported by the DFG through the Research Training Group “Molecular Principles of Synthetic Biology” (GRK 2062).

## AUTHOR CONTRIBUTIONS

EF, MSS and FCS initialized the project. EF and FCS planned the project. EF and AM performed the experiments. EF, AM and FCS wrote the manuscript and all authors discussed the results and commented on the manuscript.

## Table of contents figure

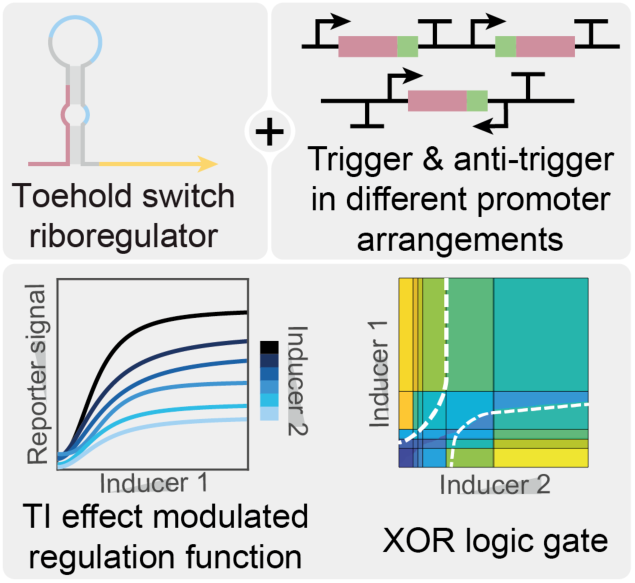

Posttranscriptional toehold switch riboregulators are used in combination with activating trigger and deactivating anti-trigger RNAs to study transcriptional interference effects within different arrangements of inducible promoters. The best performing designs were used to implement an RNA-based XOR gate.

