## Supporting Information for "Transcriptional interference in toehold switch-based RNA circuits"

### Supplementary Information

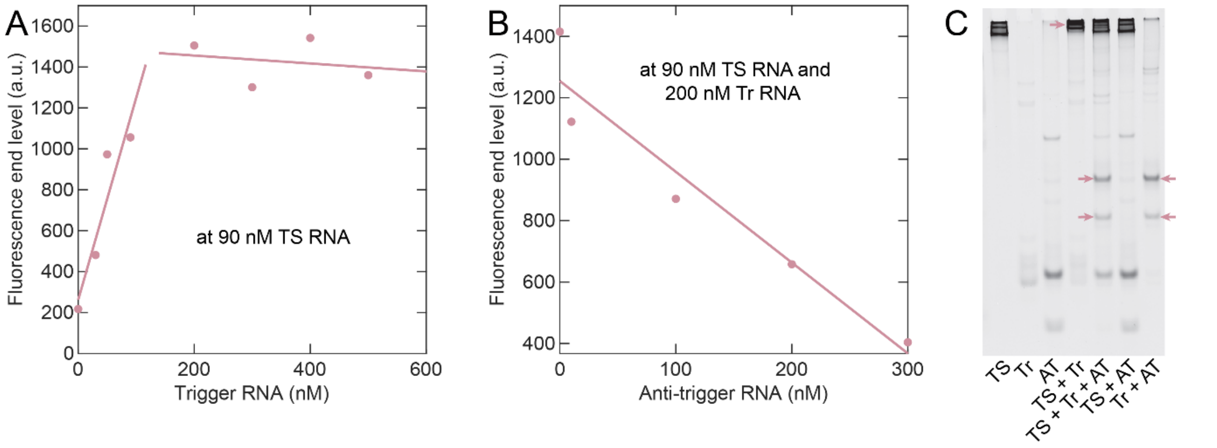


**Figure S1:** Purified toehold switch (TS), trigger (Tr) and anti-trigger (AT) RNA in a cell-free gene expression test (TXTL test in cell extract). (A) Different Tr RNA concentrations at a fix TS-CFP RNA concentration of 90 nM. Increasing amounts of Tr proportionally increase the end level, but the end level saturates for a Tr concentration of around 90 nM. (B) TXTL test for different AT RNA concentrations at a fix TS concentration of 90 nM and a fix Tr concentration of 200 nM. The end level decreases for increasing anti-trigger concentrations steadily. The linear fit crosses the x-axis at around 400 nM AT concentration. (C) Electrophoretic mobility shift assay (EMSA). TS RNA shows three bands possibly due to different secondary structures. Tr RNA shows one band and AT RNA shows tow bands corresponding to transcripts terminated after either the first or the second terminator (double terminator on template). Hybridization of TS and Tr cause a shift of the TS band (red arrow). Hybridization of Tr and AT form two new bands in presence and absence of TS. Interactions between TS and AT were not observed.

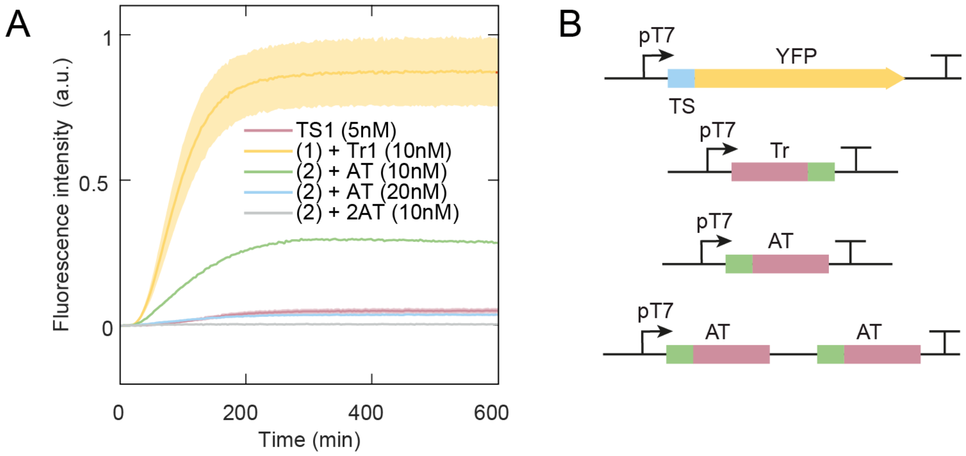


**Figure S2:** (A) TXTL test in PURExpress with linear templates coding for toehold switch, trigger or anti-trigger under the control of a T7 promoter. TS can be activated, if Tr is cotranscribed and deactivated if also AT is cotranscribed. The signal level reaches only the leaky expression level of the TS for a double excess of AT template compared to Tr template concentration. The same signal level is reached for a template coding for two AT in a row which is added in the same concentration as the trigger template. (B) Linear templates used for the experiment shown in (A).

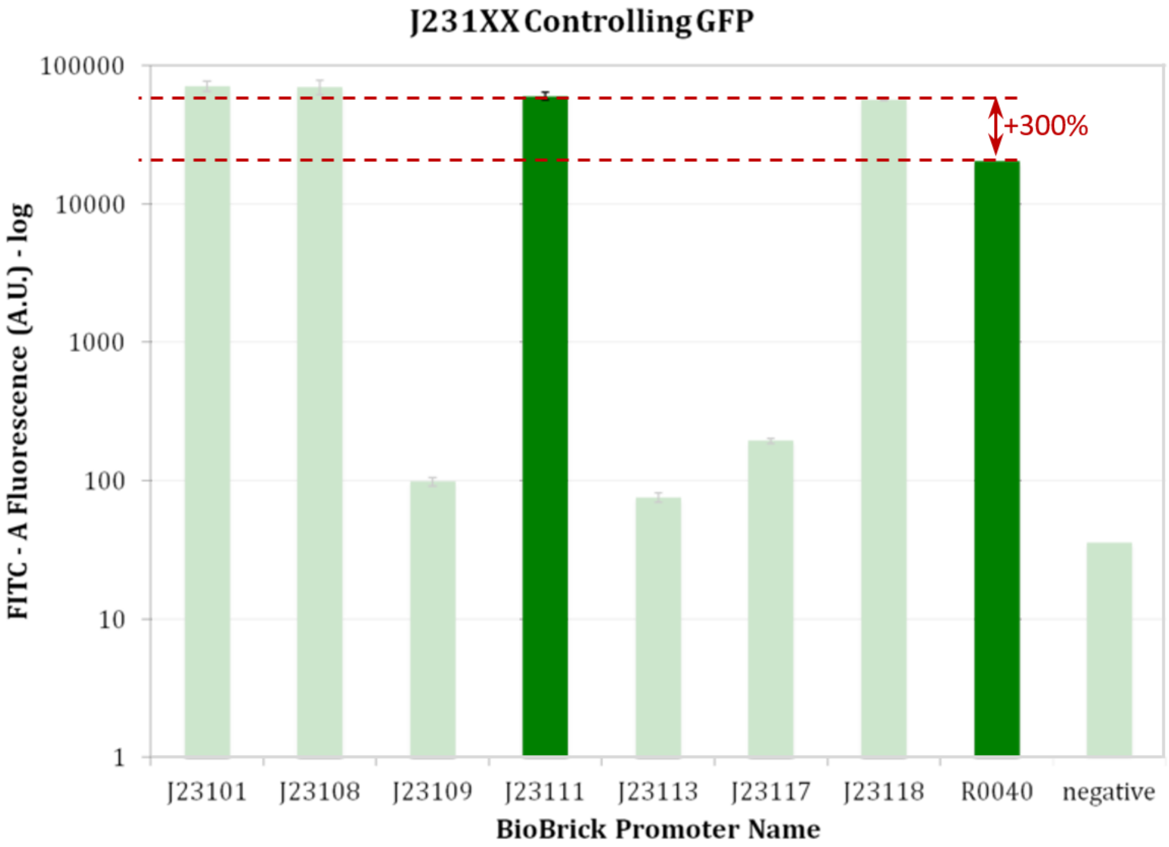


**Figure S3:** The promoter strength of J23111 is about three times higher than for pLTet-O1,[1] which is registered as R0040 in the iGEM repository. Figure was adapted of the iGEM team Boston 2012.[2]

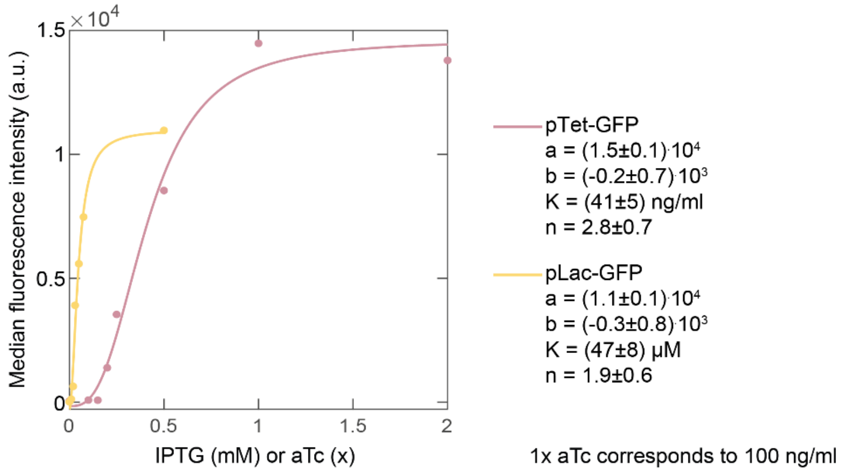


**Figure S4:** Hill curves for pTet-GFP and pLac-GFP. A plasmid containing pTet-GFP or pLac-GFP was transformed to DH5alphaZ1. Overnight cultures were diluted 1:100 and induced with up to 200 ng/ml aTc or up to 0.5 mM IPTG at OD 0.3. After overnight induction samples were measured in a flow cytometer (CyFlow Cube 6, Sysmex) and the median fluorescence signal was plotted against the inducer concentration. Hill curves were fitted to the data and the fitting parameters are shown in the plots. The Tet promoter strength is about 36% higher, which confirms the results of Bujard et al.[1]

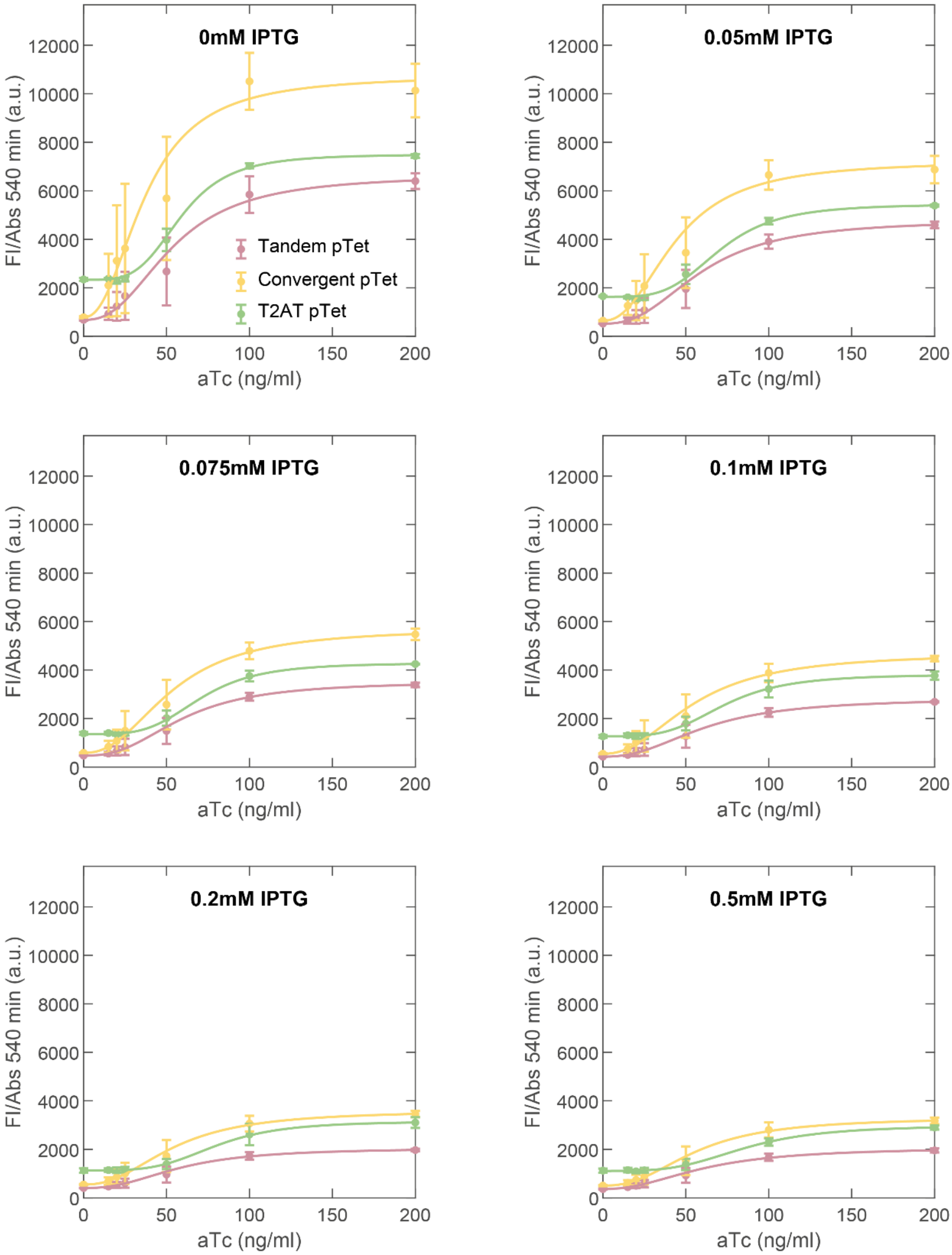


**Figure S5:** Full data set for the pTet-Tr promoter designs. (A) FI/Abs signal after 540 min at constant IPTG concentrations. Hill curves were fitted to the data. Convergent pTet shows the highest signals for 0 mM IPTG. T2AT is also upregulated at this IPTG concentration. The signals are repressed for increasing IPTG concentrations. Convergent pTet and T2AT pTet reach the same level for I0.5a200, Tandem pTet is shifted to smaller values.

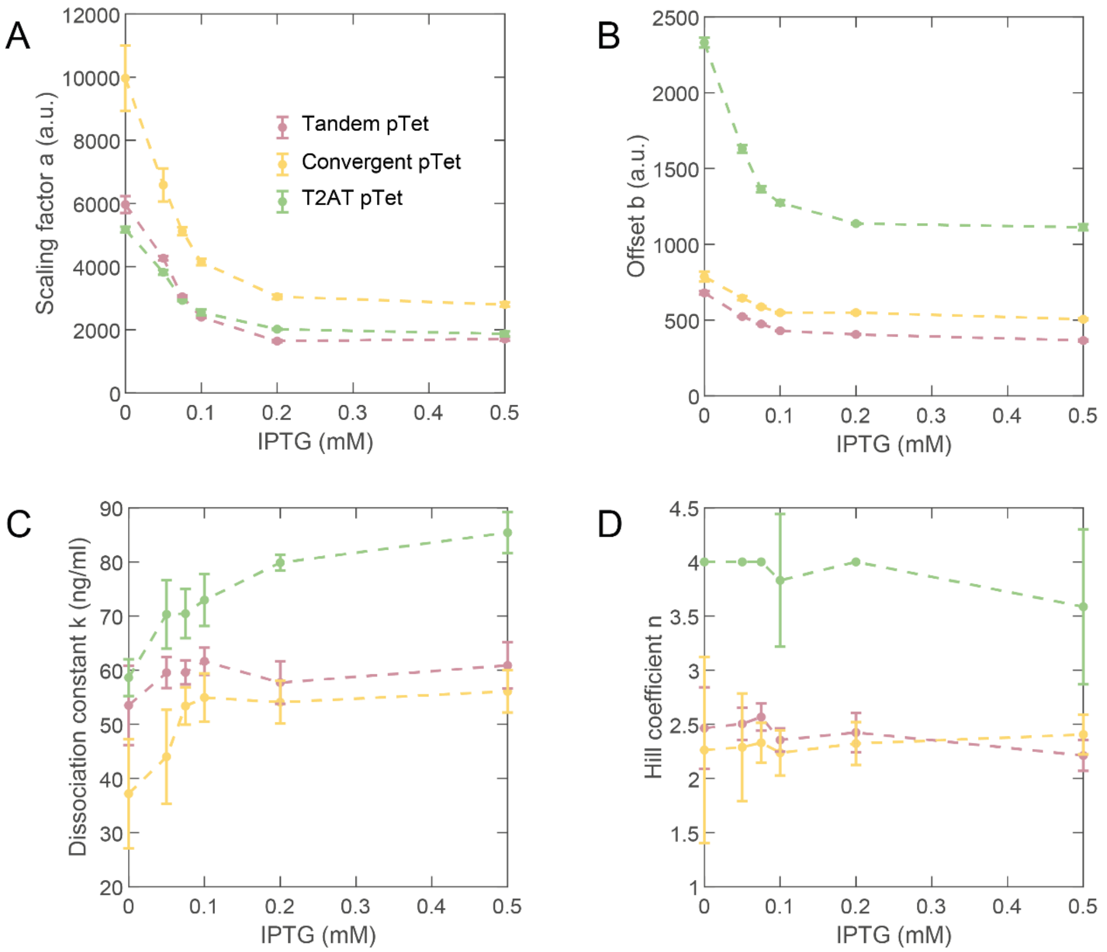


**Figure S6:** Scaling factor a, offset b, dissociation constant k and hill coefficient n depending on anti-trigger inducer concentration (IPTG) for pTet-Tr designs. The course of the offset b is discussed below, see main text for a detailed discussion of parameters a, k and n.

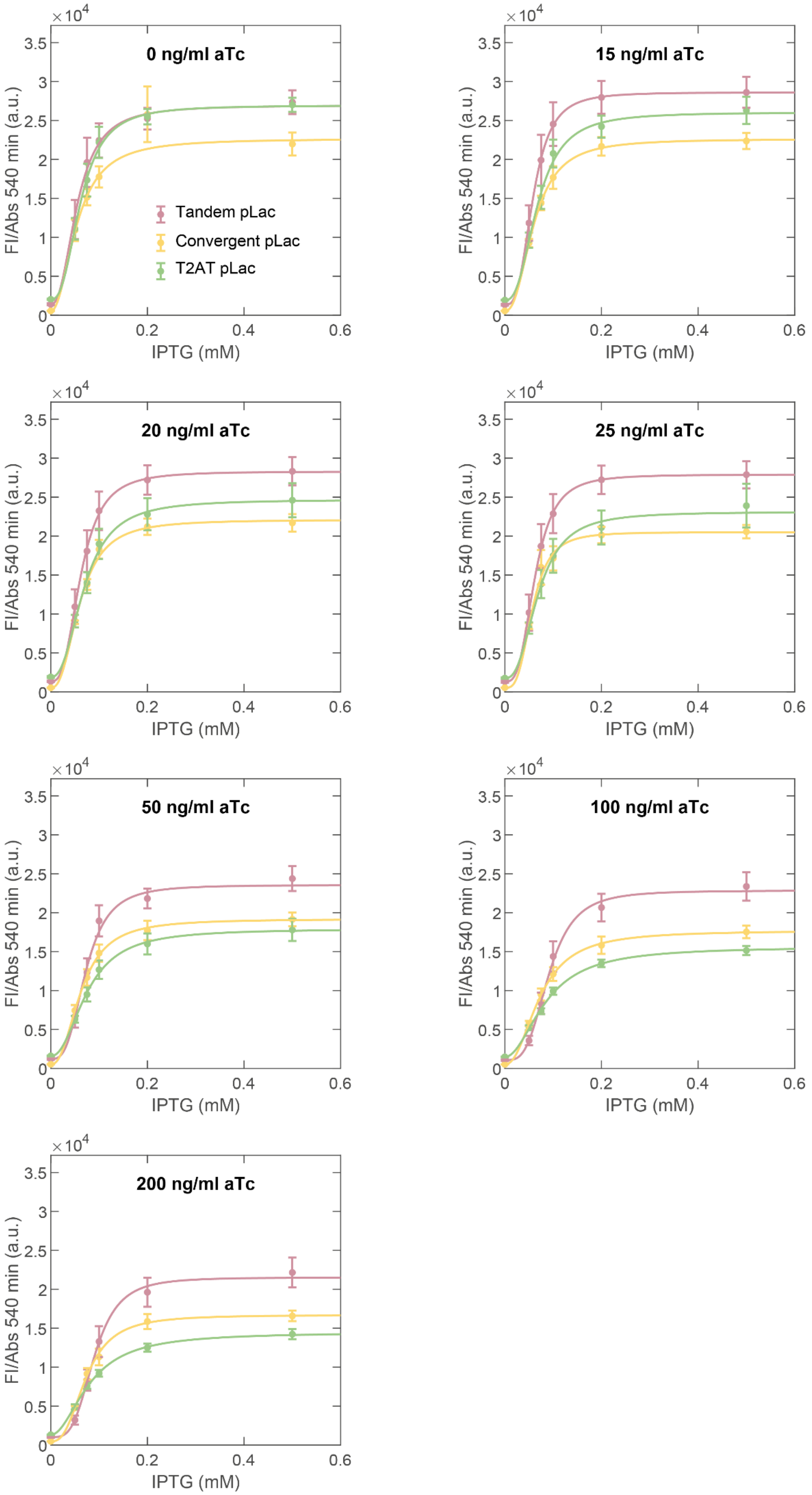
0 ng/ml aTc
15 ng/ml aTc
20 ng/ml aTc
25 ng/ml aTc
50 ng/ml aTc
100 ng/ml aTc
200 ng/ml aTc
Tandem pLac
Convergent pLac
T2AT pLac
FI/Abs 540 min (a.u.)
IPTG (mM)
×10^4
0
0.5
1
1.5
2
2.5
3
3.5
0.2
0.4
0.6

**Figure S7:** Full data set for pLac promoter designs. (A) FI/Abs signal after 540 min of induction at constant aTc concentrations. Hill curves were fitted to the data. The induction curves for 0 ng/ml aTc almost overlap for all designs. A higher signal repression under aTc induction can be observed for the convergent pLac and T2AT pLac design.

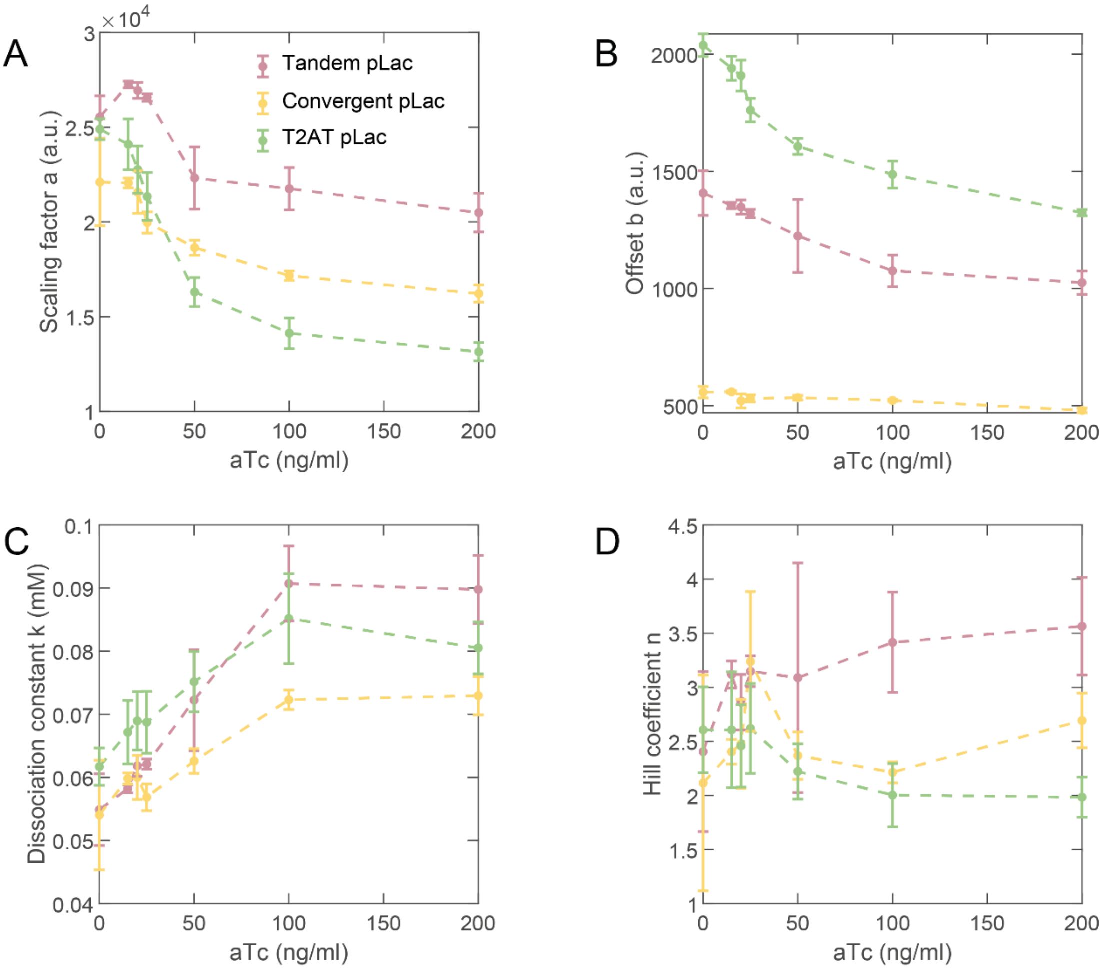


**Figure S8:** Scaling factor a, offset b, dissociation constant k and hill coefficient n depending on anti-trigger inducer concentration (aTc) for the pLac-Tr designs. The course of the offset b is discussed below, see main text for a detailed discussion of parameters a, k and n.

.

**Effects on offset b**

The parameter b gives the course of the 3D landscape in absence of trigger but under of anti-trigger induction. The offset is for all designs except T2AT pTet small compared to the ON signals. All designs show a signal decrease for increasing AT concentrations, but the steepest decrease was observed for the T2AT designs.

The offset in the tandem pLac design is more than 2 fold higher compared to the tandem pTet design. Convergent pTet shows the same offset than its reference systems. The convergent pLac design shows the lowest offset of all systems. As we observed no TS-AT hybridization in the EMSA study (Figure S1), we expect no direct effect of AT on TS. The signal decrease for the tandem designs might be a result of leaky transcription of the promoters and a small background concentration of Tr, which can be compensated by AT and explains a signal decrease. For the convergent and T2AT designs also dislodgement events by AT transcribing RNAPs can increase the background Tr concentration, explaining a higher signal decrease with increasing AT concentrations.

As we already observed an upregulating effect on J23111 due to a close pTet promoter, we also expect a higher offset in the Tandem design compared to the Tandem pTet design (Figure S6 and S8).

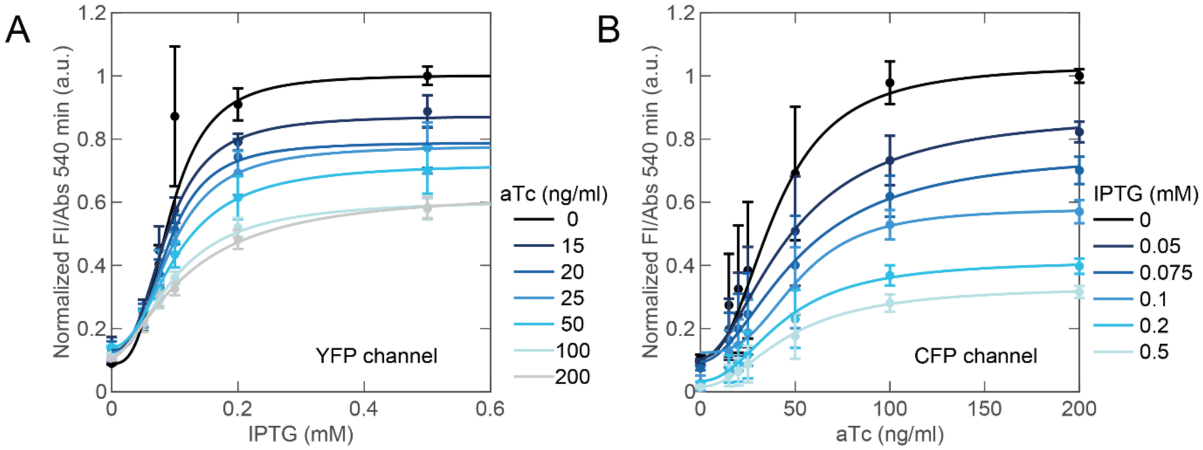


**Figure S9:** A XOR logic gate was assembled by combining a T2AT pLac design including a YFP reporter (XOR YFP) and a convergent pTet design including a CFP reporter (XOR CFP). The FI/Abs signals after 540 min of induction were normalized to the signal at I0.5a0 or I0a200 respectively and Hill curves were fitted. XOR YFP (A) shows a higher offset compared to XOR CFP (B), aTc induction causes a signal decrease down to 60% of the maximum (compare I0.5a0 and I0.5a200). XOR CFP can be repressed to less than 40% of the maximum under addition of IPTG (compare I0a200 and I0.5a200).

**Figure S10:** Hill fit parameters a, b, k and n for the XOR gates

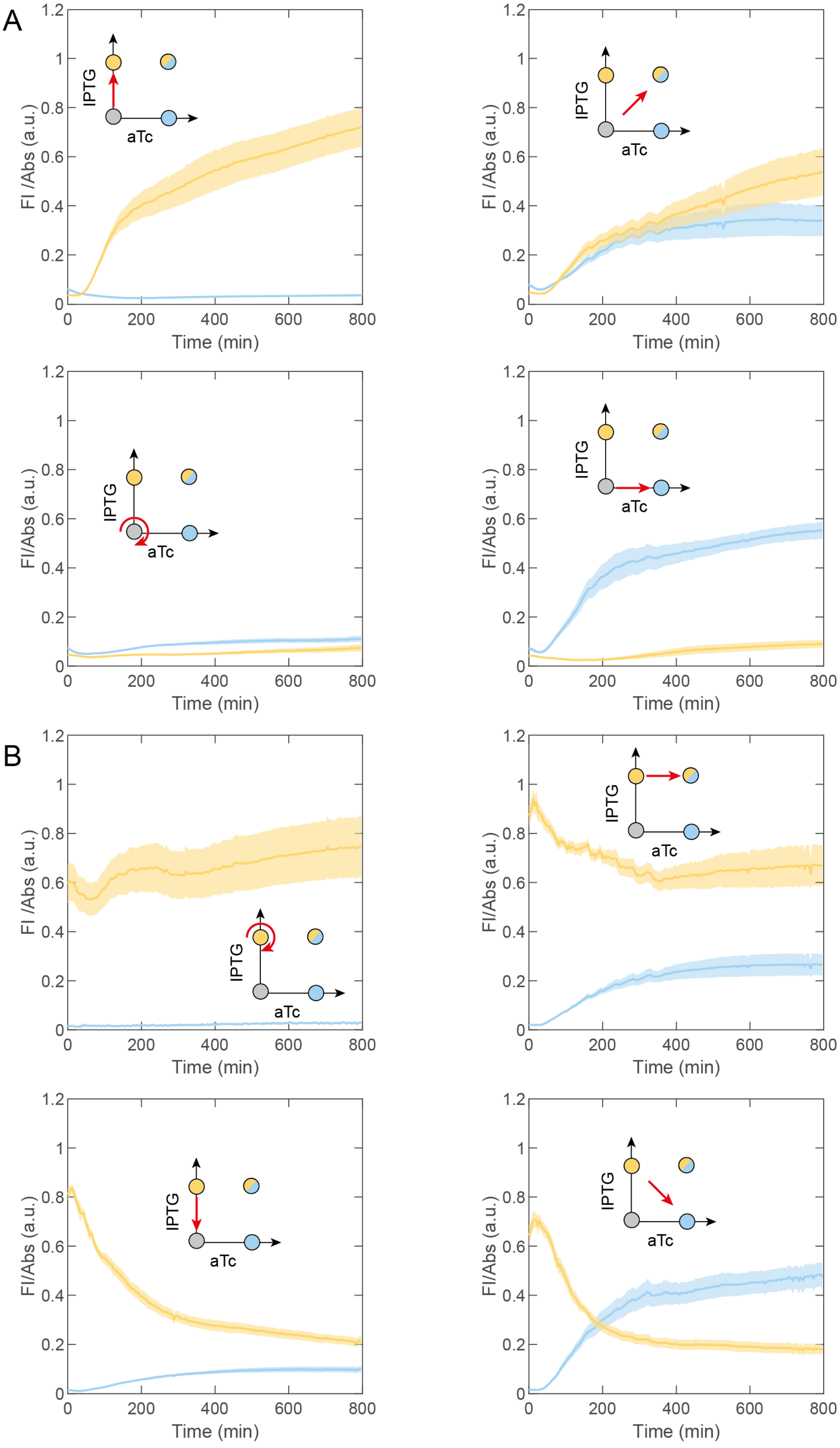
A
B
FI /Abs (a.u.)
FI/Abs (a.u.)
Time (min)
IPTG
aTc
0
0.2
0.4
0.6
0.8
1
1.2
200
400
600
800

**Figure S11:** XOR switching experiment. Four inducer combinations (I0a0, I1a0, I0a100 and I1a100) were chosen as the points of interest and each switching option between the points was tested. The pictograms indicate the switching path between the inducer combinations. (A) XOR switching experiment with I0a0 as the starting point. Both inducers stay at a low level, if the inducer concentrations are kept constant. XOR gate shows a signal increase for one of the reporters at a low level of the second reporter, if just one inducer is added. If both inducers are added, both reporters are expressed, but the signal level is reduced compared to the fully induced states (intermediate state). (B) XOR switching with I1a0 as the starting point. The YFP signal level stays high and the CFP signal stays low, if the inducer conditions are kept constant. The YFP signal decreases for all other inducer combinations to either an intermediate state (I1a100) or a low level (I0a0, I0a100). Low CFP levels are observed in absence of aTc. CFP level increases to an intermediate state, if both inducers are added and to a high state, if just aTc and no IPTG is added.

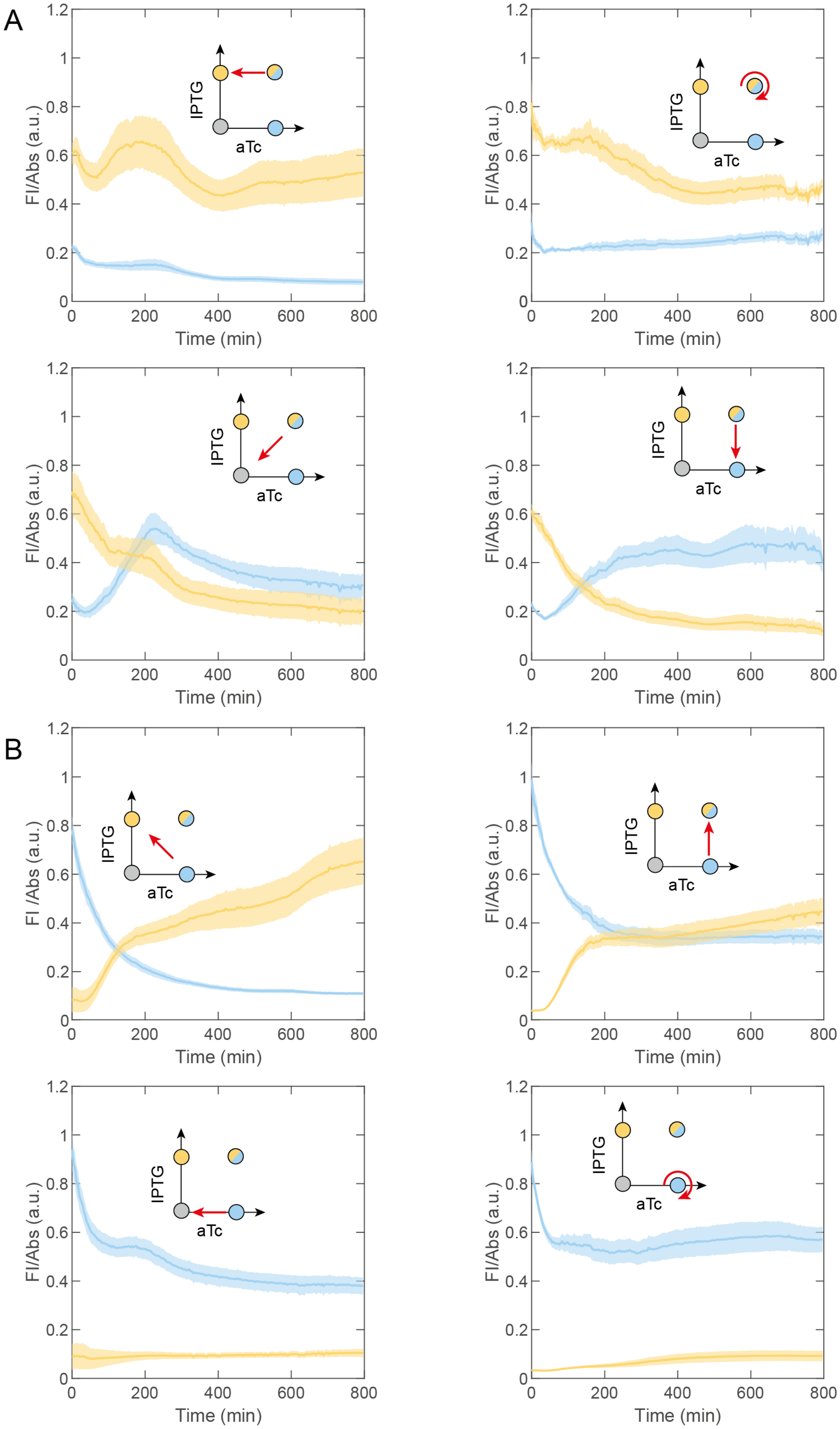
A
B
FI/Abs (a.u.)
FI /Abs (a.u.)
Time (min)
IPTG
aTc
0
0.2
0.4
0.6
0.8
1
1.2
200
400
600
800

**Figure S12:** XOR switching experiment. (A) XOR switching with I1a100 as starting point. The switching behavior matches again the expectations shown in the pictograms. (B) Switching experiment with I0a100 as starting point. A CFP signal decrease is observed for all switching experiments down to an intermediate or a low state. The YFP signal level increases to an intermediate (both inducers are present) or high state (no aTc, but IPTG is present).

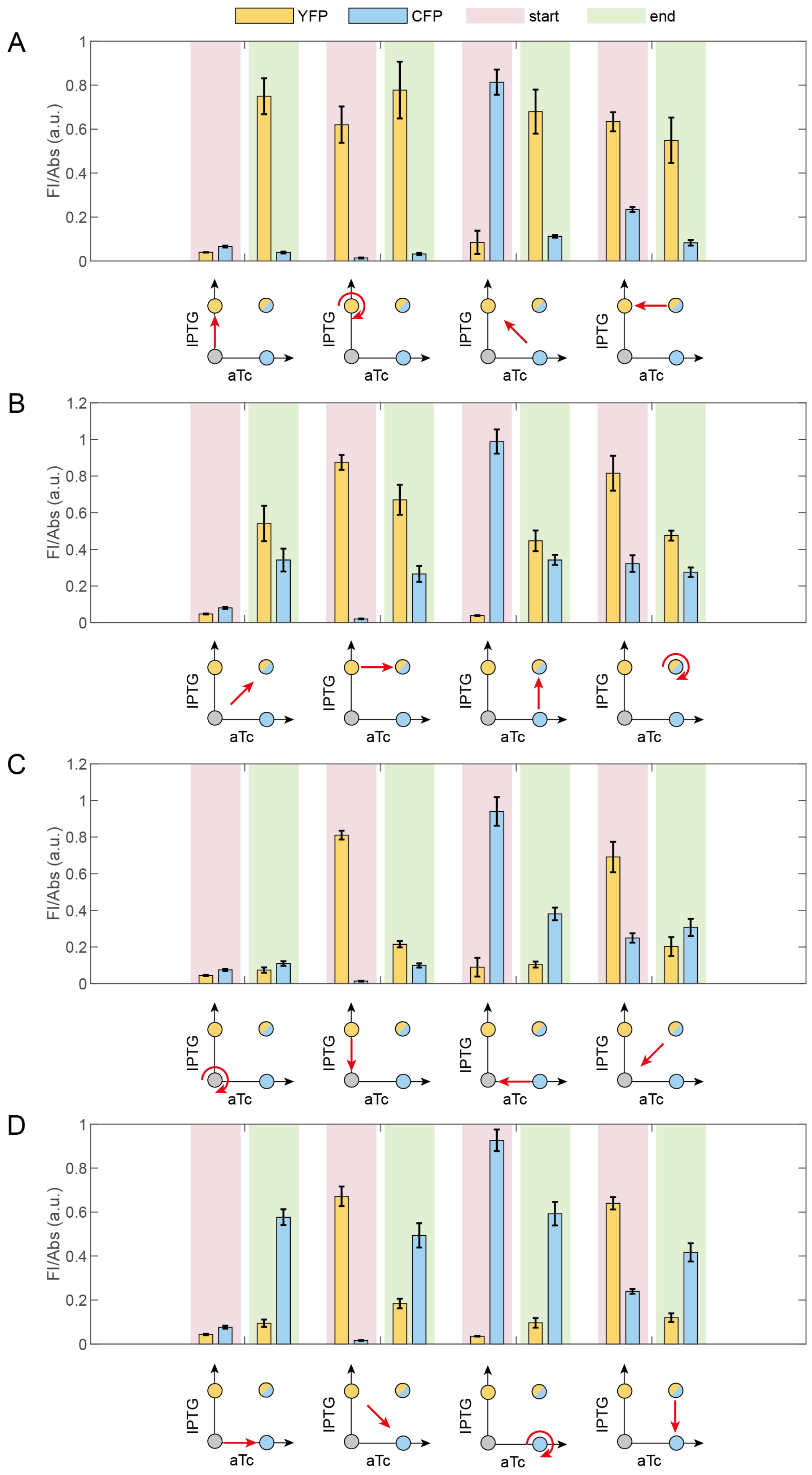


**Figure S13:** The results of Figure 8 and 9 are summarized in bar plots showing the start and end levels of the XOR reporters. The pictograms indicate the switching path between the inducer combinations I0a0, I1a0, I0a100 and I1a100. Starting from different positions, the end position is always the same within the bar graphs A-D and the end levels match qualitatively.

**Supplementary Table 1:** ON/OFF ratios for all designs. ON state is defined as the signal level at maximum trigger inducer concentration in absence of anti-trigger inducer. The OFF state is defined as the signal level with both inducers at their maximum concentration or as signal level in absence of any inducers. The ON/OFF ratios were calculated after 540 min of induction.

| Design | ON/OFF ratio (I0+a200/I0.5+a200) | ON/OFF ratio (I0+a200/I0+a0) |
|---|---|---|
| Tandem pTet | 3.3±0.2 | 9.4±0.6 |
| convergent pTet | 3.2±0.4 | 13±2 |
| T2AT pTet | 2.6±0.1 | 3.2±0.1 |
| XOR CFP | 3.2±0.2 | 9±1 |
| **Design** | **ON/OFF ratio** (I0.5+a0/I0.5+a200) | **ON/OFF ratio** (I0.5+a0/I0+a0) |
| Tandem pLac | 1.2±0.1 | 19±2 |
| Convergent pLac | 1.3±0.1 | 39±3 |
| T2AT pLac | 1.9±0.1 | 13.3±0.7 |
| XOR YFP | 1.7±0.1 | 11.2±0.5 |

**Supplementary Table 2:** pSB1A3-Tandem promoters (pLac-Tr)

```
gacgtcataaatgtgagcggataacattgacattgtgagcggataacaagatactgagcacatcaactgactattctgtg
caatagacagtaaagcagggataaacgagatagataagataagaatcctaccatgatccaggcatcaaataaaacgaaag
gctcagtcgaaagactgggcctttcgttttatctgttgtttgtcggtgaacgctctctactagagtcacactggctcacc
ttcgggtgggcctttctgcgtttatatgatgcgtagggagatccggaatctattggcctatgtcacctcgagtccctatc
agtgatagagattgacatccctatcagtgatagagatactgagcacatcatggtaggattcttatcttatctatctcgtt
tatccctgctttactgtctattgcacagaatagtcagtggacgcaaaaaaccccgcttcggcggggtttttttcgcgtgcg
ctgatctgagttgacggctagctcagtcctaggtatagtgctagctcttatcttatctatctcgtttatccctgcataca
gaaacagaggagatacgcaatgataaacgagaacctggcggcagcgcaaaagatgagcaaaggcgaagaactgttcacgg
gtgtggttccgatcctggttgaactggatggcgatgtgaacggtcataaatttagcgtgtctggtgaaggcgaaggtgat
gcgacctacggcaaactgacgctgaaactgatttgcaccacgggtaaactgccggttccgtggccgaccctggtgaccac
gctgggttatggtctgatgtgtttcgcacgttacccggatcacatgaaacgccatgatttctttaaatctgcgatgccgg
aaggctatgtgcaggaacgtaccatctttttcaaagatgatggtaactacaaaacccgcgcggaagttaaatttgaaggc
gatacgctggtgaaccgtattgaactgaaaggtatcgatttcaaagaagatggcaatattctgggtcacaaactggaata
caactacaacagtcataacgtgtacattaccgccgataaacagaaaaacggtatcaaagcaaacttcaaaatccgtcaca
acatcgaagatggcggtgttcagctggccgatcattaccagcagaacaccccgattggcgatggtccggtgctgctgccg
gataatcattatctgagttaccagagcaaactgtctaaagatccgaatgaaaaacgcgatcacatggttctgctggaatt
tgtgaccgcggccggcattacgcatggtatggatgaactgtataaataaccaggcatcaaataaaacgaaaggctcagtc
gaaagactgggcctttcgttttatctgttgtttgtcggtgaacgctctctactagagtcacactggctcaccttcgggtg
ggcctttctgcgtttataactagtgcgctgcagtccggcaaaaaagggcaaggtgtcaccaccctgcccttttttcttttaa
aaccgaaaagattacttcgcgttatgcaggcttcctcgctcactgactcgctgcgctcggtcgttcggctgcggcgagcg
gtatcagctcactcaaaggcggtaatacggttatccacagaatcaggggataacgcaggaaagaacatgtgagcaaaagg
ccagcaaaaggccaggaaccgtaaaaaggccgcgttgctggcgtttttccacaggctccgcccccctgacgagcatcaca
aaaatcgacgctcaagtcagaggtggcgaaacccgacaggactataaagataccaggcgtttccccctggaagctccctc
gtgcgctctcctgttccgaccctgccgcttaccggatacctgtccgcctttctcccttcgggaagcgtggcgctttctca
tagctcacgctgtaggtatctcagttcggtgtaggtcgttcgctccaagctgggctgtgtgcacgaaccccccgttcagc
ccgaccgctgcgccttatccggtaactatcgtcttgagtccaacccggtaagacacgacttatcgccactggcagcagcc
actggtaacaggattagcagagcgaggtatgtaggcggtgctacagagttcttgaagtggtggcctaactacggctacac
tagaagaacagtatttggtatctgcgctctgctgaagccagttaccttcggaaaaagagttggtagctcttgatccggca
aacaaaccaccgctggtagcggtggtttttttgtttgcaagcagcagattacgcgcagaaaaaaaggatctcaagaagat
cctttgatcttttctacggggtctgacgctcagtggaacgaaaactcacgttaagggattttggtcatgagattatcaaa
aaggatcttcacctagatccttttaaattaaaaatgaagttttaaatcaatctaaagtatatatgagtaaacttggtctg
acagttaccaatgcttaatcagtgaggcacctatctcagcgatctgtctatttcgttcatccatagttgcctgactcccc
gtcgtgtagataactacgatacgggagggcttaccatctggccccagtgctgcaatgataccgcgagacccacgctcacc
ggctccagatttatcagcaataaaccagccagccggaagggccgagcgcagaagtggtcctgcaactttatccgcctcca
tccagtctattaattgttgccgggaagctagagtaagtagttcgccagttaatagtttgcgcaacgttgttgccattgct
acaggcatcgtggtgtcacgctcgtcgtttggtatggcttcattcagctccggttcccaacgatcaaggcgagttacatg
atcccccatgttgtgcaaaaaagcggttagctccttcggtcctccgatcgttgtcagaagtaagttggccgcagtgttat
cactcatggttatggcagcactgcataattctcttactgtcatgccatccgtaagatgcttttctgtgactggtgagtac
tcaaccaagtcattctgagaatagtgtatgcggcgaccgagttgctcttgcccggcgtcaatacgggataataccgcgcc
acatagcagaactttaaaagtgctcatcattggaaaacgttcttcggggcgaaaactctcaaggatcttaccgctgttga
gatccagttcgatataacccactcgtgcacccaactgatcttcagcatcttttactttcaccagcgtttctgggtgagca
aaaacaggaaggcaaaatgccgcaaaaaagggaataagggcgacacggaaatgttgaatactcatactcttcctttttca
atattattgaagcatttatcagggttattgtctcatgagcggatacatatttgaatgtatttagaaaaataaacaaatag
gggttccgcgcacatttccccgaaaagtgccacct
```

**Supplementary Table 3:** pSB1A3-convergent promoters (pLac-Tr)

cgcgaaaaaaccccgccgaagcggggtttttttgcgataaatgtgagcggataacattgacattgtgagcggataacaaga
tactgagcacatccactgactattctgtgcaatagacagtaaagcagggataaacgagatagataagataagaatggaac
catgatgtgctcagtatctctatcactgatagggatgtcaatctctatcactgatagggaccaggcatcaaataaaacga
aaggctcagtcgaaagactgggcctttcgttttatctgttgtttgtcggtgaacgctctctactagagtcacactggctc
accttcgggtgggcctttctgcgtttatagaattcttgacggctagctcagtcctaggtatagtgctagcgggtcttatc
ttatctatctcgtttatccctgcatacagaaacagaggagatacgcaatgataaacgagaacctggcggcagcgcaaaag
atgagcaaaggcgaagaactgttcacgggtgtggttccgatcctggttgaactggatggcgatgtgaacggtcataaatt
tagcgtgtctggtgaaggcgaaggtgatgcgacctacggcaaactgacgctgaaactgatttgcaccacgggtaaactgc
cggttccgtggccgaccctggtgaccacgctgggttatggtctgatgtgtttcgcacgttacccggatcacatgaaacgc
catgatttctttaaatctgcgatgccggaaggctatgtgcaggaacgtaccatctttttcaaagatgatggtaactacaa
aacccgcgcggaagttaaatttgaaggcgatacgctggtgaaccgtattgaactgaaaggtatcgatttcaaagaagatg
gcaatattctgggtcacaaactggaatacaactacaacagtcataacgtgtacattaccgccgataaacagaaaaacggt
atcaaagcaaacttcaaaatccgtcacaacatcgaagatggcggtgttcagctggccgatcattaccagcagaacacccc
gattggcgatggtccggtgctgctgccggataatcattatctgagttaccagagcaaactgtctaaagatccgaatgaaa
aacgcgatcacatggttctgctggaatttgtgaccgcggccggcattacgcatggtatggatgaactgtataaataacca
ggcatcaaataaaacgaaaggctcagtcgaaagactgggcctttcgttttatctgttgtttgtcggtgaacgctctctac
tagagtcacactggctcaccttcgggtgggcctttctgcgtttataactagtgcgctgcagtccggcaaaaaagggcaag
gtgtcaccaccctgccctttttctttaaaaccgaaaagattacttcgcgttatgcaggcttcctcgctcactgactcgct
gcgctcggtcgttcggctgcggcgagcggtatcagctcactcaaaggcggtaatacggttatccacagaatcaggggata
acgcaggaaagaacatgtgagcaaaaggccagcaaaaggccaggaaccgtaaaaaggccgcgttgctggcgtttttccac
aggctccgcccccctgacgagcatcacaaaaatcgacgctcaagtcagaggtggcgaaacccgacaggactataaagata
ccaggcgtttccccctggaagctccctcgtgcgctctcctgttccgaccctgccgcttaccggatacctgtccgcctttc
tcccttcgggaagcgtggcgctttctcatagctcacgctgtaggtatctcagttcggtgtaggtcgttcgctccaagctg
ggctgtgtgcacgaaccccccgttcagcccgaccgctgcgccttatccggtaactatcgtcttgagtccaacccggtaag
acacgacttatcgccactggcagcagccactggtaacaggattagcagagcgaggtatgtaggcggtgctacagagttct
tgaagtggtggcctaactacggctacactagaagaacagtatttggtatctgcgctctgctgaagccagttaccttcgga
aaaagagttggtagctcttgatccggcaaacaaaccaccgctggtagcggtggtttttttgtttgcaagcagcagattac
gcgcagaaaaaaaggatctcaagaagatcctttgatcttttctacggggtctgacgctcagtggaacgaaaactcacgtt
aagggattttggtcatgagattatcaaaaaggatcttcacctagatccttttaaattaaaaatgaagttttaaatcaatc
taaagtatatatgagtaaacttggtctgacagttaccaatgcttaatcagtgaggcacctatctcagcgatctgtctatt
tcgttcatccatagttgcctgactccccgtcgtgtagataactacgatacgggagggcttaccatctggccccagtgctg
caatgataccgcgagacccacgctcaccggctccagatttatcagcaataaaccagccagccggaagggccgagcgcaga
agtggtcctgcaactttatccgcctccatccagtctattaattgttgccgggaagctagagtaagtagttcgccagttaa
tagtttgcgcaacgttgttgccattgctacaggcatcgtggtgtcacgctcgtcgtttggtatggcttcattcagctccg
gttcccaacgatcaaggcgagttacatgatcccccatgttgtgcaaaaaagcggttagctccttcggtcctccgatcgtt
gtcagaagtaagttggccgcagtgttatcactcatggttatggcagcactgcataattctcttactgtcatgccatccgt
aagatgcttttctgtgactggtgagtactcaaccaagtcattctgagaatagtgtatgcggcgaccgagttgctcttgcc
cggcgtcaatacgggataataccgcgccacatagcagaactttaaaagtgctcatcattggaaaacgttcttcggggcga
aaactctcaaggatcttaccgctgttgagatccagttcgatataacccactcgtgcacccaactgatcttcagcatcttt
tactttcaccagcgtttctgggtgagcaaaaacaggaaggcaaaatgccgcaaaaaagggaataagggcgacacggaaat
gttgaatactcatactcttcctttttcaatattattgaagcatttatcagggttattgtctcatgagcggatacatattt
gaatgtatttagaaaaataaacaaataggggttccgcgcacatttccccgaaaagtgccacctgacgt

**Supplementary Table 4:** pSB1A3-T2AT (pLac-Tr)

actcttcctttttcaatattattgaagcatttatcagggttattgtctcatgagcggatacatatttgaatgtatttaga
aaaataaacaaataggggttccgcgcacatttccccgaaaagtgccacctgacgtctaagaagacctcgcgaaaaaaccc
cgccgaagcggggtttttttgcgcgaccacgacggctaaaaaaaaccccgccctgtcaggggcggggtttttttttgagcg
ggactgactattctgtgcaatagtcagtaaagcagggataaacgagatagataagataagatagatcctaccatgattta
aacaaaattatttgtagaggactgtttcggcccttttgggccatcgtcaggtcggatacacatccggcgacagtctctcg
agataaatgtgagcggataacattgacattgtgagcggataacaagatactgagcacagggactgactattctgtgcaat
agtcagtaaagcagggataaacgagatagataagataagatagatcctaccatgatgtgctcagtatctctatcactgat
agggatgtcaatctctatcactgatagggagagaccaggcatcaaataaaacgaaaggctcagtcgaaagactgggcctt
tcgttttatctgttgtttgtcggtgaacgctctctactagagtcacactggctcaccttcgggtgggcctttctgcgttt
atagtgcgctgacaagtccattaccaaggagtgcgatgcagttactcagaattcttgacggctagctcagtcctaggtat
agtgctagcgggtcttatcttatctatctcgtttatccctgcatacagaaacagaggagatacgcaatgataaacgagaa
cctggcggcagcgcaaaagatgagcaaaggcgaagaactgttcacgggtgtggttccgatcctggttgaactggatggcg
atgtgaacggtcataaatttagcgtgtctggtgaaggcgaaggtgatgcgacctacggcaaactgacgctgaaactgatt
tgcaccacgggtaaactgccggttccgtggccgaccctggtgaccacgctgggttatggtctgatgtgtttcgcacgtta
cccggatcacatgaaacgccatgatttctttaaatctgcgatgccggaaggctatgtgcaggaacgtaccatctttttca
aagatgatggtaactacaaaacccgcgcggaagttaaatttgaaggcgatacgctggtgaaccgtattgaactgaaaggt
atcgatttcaaagaagatggcaatattctgggtcacaaactggaatacaactacaacagtcataacgtgtacattaccgc
cgataaacagaaaaacggtatcaaagcaaacttcaaaatccgtcacaacatcgaagatggcggtgttcagctggccgatc
attaccagcagaacaccccgattggcgatggtccggtgctgctgccggataatcattatctgagttaccagagcaaactg
tctaaagatccgaatgaaaaacgcgatcacatggttctgctggaatttgtgaccgcggccggcattacgcatggtatgga
tgaactgtataaataaccaggcatcaaataaaacgaaaggctcagtcgaaagactgggcctttcgttttatctgttgttt

```
gtcggtgaacgctctctactagagtcacactggctcaccttcgggtgggcctttctgcgtttataactagtgcgctgcag
tccggcaaaaaagggcaaggtgtcaccaccctgccctttttctttaaaaccgaaaagattacttcgcgttatgcaggctt
cctcgctcactgactcgctgcgctcggtcgttcggctgcggcgagcggtatcagctcactcaaaggcggtaatacggtta
tccacagaatcaggggataacgcaggaaagaacatgtgagcaaaaggccagcaaaaggccaggaaccgtaaaaaggccgc
gttgctggcgtttttccacaggctccgcccccctgacgagcatcacaaaaatcgacgctcaagtcagaggtggcgaaacc
cgacaggactataaagataccaggcgtttccccctggaagctccctcgtgcgctctcctgttccgaccctgccgcttacc
ggatacctgtccgcctttctcccttcgggaagcgtggcgctttctcatagctcacgctgtaggtatctcagttcggtgta
ggtcgttcgctccaagctgggctgtgtgcacgaaccccccgttcagcccgaccgctgcgccttatccggtaactatcgtc
ttgagtccaacccggtaagacacgacttatcgccactggcagcagccactggtaacaggattagcagagcgaggtatgta
ggcggtgctacagagttcttgaagtggtggcctaactacggctacactagaagaacagtatttggtatctgcgctctgct
gaagccagttaccttcggaaaaagagttggtagctcttgatccggcaaacaaaccaccgctggtagcggtggtttttttg
tttgcaagcagcagattacgcgcagaaaaaaaggatctcaagaagatcctttgatcttttctacggggtctgacgctcag
tggaacgaaaactcacgttaagggattttggtcatgagattatcaaaaaggatcttcacctagatccttttaaattaaaa
atgaagttttaaatcaatctaaagtatatatgagtaaacttggtctgacagttaccaatgcttaatcagtgaggcaccta
tctcagcgatctgtctatttcgttcatccatagttgcctgactccccgtcgtgtagataactacgatacgggagggctta
ccatctggccccagtgctgcaatgataccgcgagacccacgctcaccggctccagatttatcagcaataaaccagccagc
cggaagggccgagcgcagaagtggtcctgcaactttatccgcctccatccagtctattaattgttgccgggaagctagag
taagtagttcgccagttaatagtttgcgcaacgttgttgccattgctacaggcatcgtggtgtcacgctcgtcgtttggt
atggcttcattcagctccggttcccaacgatcaaggcgagttacatgatcccccatgttgtgcaaaaaagcggttagctc
cttcggtcctccgatcgttgtcagaagtaagttggccgcagtgttatcactcatggttatggcagcactgcataattctc
ttactgtcatgccatccgtaagatgcttttctgtgactggtgagtactcaaccaagtcattctgagaatagtgtatgcgg
cgaccgagttgctcttgcccggcgtcaatacgggataataccgcgccacatagcagaactttaaaagtgctcatcattgg
aaaacgttcttcggggcgaaaactctcaaggatcttaccgctgttgagatccagttcgatataacccactcgtgcaccca
actgatcttcagcatcttttactttcaccagcgtttctgggtgagcaaaaacaggaaggcaaaatgccgcaaaaaaggga
ataagggcgacacggaaatgttgaatactcat
```

**Supplementary Table 5:** pSB1A3-Tandem pTet-Tr

```
tgaatactcatactcttcctttttcaatattattgaagcatttatcagggttattgtctcatgagcggatacatatttga
atgtatttagaaaaataaacaaataggggttccgcgcacatttccccgaaaagtgccacctgacgtctaagaagacttct
ccctatcagtgatagagattgacatccctatcagtgatagagatactgagcacatcagggactgactattctgtgcaata
gtcagtaaagcagggataaacgagatagataagataagatagatcctaccatgatccaggcatcaaataaaacgaaaggc
tcagtcgaaagactgggcctttcgttttatctgttgtttgtcggtgaacgctctctactagagtcacactggctcacctt
cgggtgggcctttctgcgtttatatgatgcgtagggagatcctttaggcggtgcttactcttacataaagggggctgttag
tattaccccgcgaggattacatgcctatatttgtctctttgccggcttatatggacaagcatagcattcgaaaaggtgag
ccggaaacatatgctgtctctggcacgttctattggcctatgtcacctcgagataaatgtgagcggataacattgacatt
gtgagcggataacaagatactgagcacaatcatggtaggatctatcttatcttatctatctcgtttatccctgctttact
gactattgcacagaatagtcagtcccggacgcaaaaaaccccgcttcggcggggtttttttcgctcgtactgctagtagag
gcaaaaaaaaacccgcccctgacagggcggggttttttttagccgtcgtggtgaagtccgtacacggtatgatcggacg
cctcgtgagatcaatacgtataccaggtgtcctgtgagcagcgaaagactcacaggctccgtaatactagcatgcgataa
gtccctaactgactatggccttctgattgatgccacggcgatctccgtgcgctgatctgaggaattcttgacggctagct
cagtcctaggtatagtgctagcgggtcttatcttatctatctcgtttatccctgcatacagaaacagaggagatacgcaa
tgataaacgagaacctggcggcagcgcaaaagatgagcaaaggcgaagaactgttcacgggtgtggttccgatcctggtt
gaactggatggcgatgtgaacggtcataaatttagcgtgtctggtgaaggcgaaggtgatgcgacctacggcaaactgac
gctgaaactgatttgcaccacgggtaaactgccggttccgtggccgaccctggtgaccacgctgggttatggtctgatgt
gtttcgcacgttacccggatcacatgaaacgccatgatttctttaaatctgcgatgccggaaggctatgtgcaggaacgt
accatctttttcaaagatgatggtaactacaaaacccgcgcggaagttaaatttgaaggcgatacgctggtgaaccgtat
tgaactgaaaggtatcgatttcaaagaagatggcaatattctgggtcacaaactggaatacaactacaacagtcataacg
tgtacattaccgccgataaacagaaaaacggtatcaaagcaaacttcaaaatccgtcacaacatcgaagatggcggtgtt
cagctggccgatcattaccagcagaacaccccgattggcgatggtccggtgctgctgccggataatcattatctgagtta
ccagagcaaactgtctaaagatccgaatgaaaaacgcgatcacatggttctgctggaatttgtgaccgcggccggcatta
cgcatggtatggatgaactgtataaataaccaggcatcaaataaaacgaaaggctcagtcgaaagactgggcctttcgtt
ttatctgttgtttgtcggtgaacgctctctactagagtcacactggctcaccttcgggtgggcctttctgcgtttataac
tagtgcgctgcagtccggcaaaaaagggcaaggtgtcaccaccctgccctttttctttaaaaccgaaaagattacttcgc
gttatgcaggcttcctcgctcactgactcgctgcgctcggtcgttcggctgcggcgagcggtatcagctcactcaaaggc
ggtaatacggttatccacagaatcaggggataacgcaggaaagaacatgtgagcaaaaggccagcaaaaggccaggaacc
gtaaaaaggccgcgttgctggcgtttttccacaggctccgcccccctgacgagcatcacaaaaatcgacgctcaagtcag
aggtggcgaaacccgacaggactataaagataccaggcgtttccccctggaagctccctcgtgcgctctcctgttccgac
cctgccgcttaccggatacctgtccgcctttctcccttcgggaagcgtggcgctttctcatagctcacgctgtaggtatc
tcagttcggtgtaggtcgttcgctccaagctgggctgtgtgcacgaaccccccgttcagcccgaccgctgcgccttatcc
ggtaactatcgtcttgagtccaacccggtaagacacgacttatcgccactggcagcagccactggtaacaggattagcag
agcgaggtatgtaggcggtgctacagagttcttgaagtggtggcctaactacggctacactagaagaacagtatttggta
tctgcgctctgctgaagccagttaccttcggaaaaagagttggtagctcttgatccggcaaacaaaccaccgctggtagc
ggtggtttttttgtttgcaagcagcagattacgcgcagaaaaaaaggatctcaagaagatcctttgatcttttctacggg
gtctgacgctcagtggaacgaaaactcacgttaagggattttggtcatgagattatcaaaaaggatcttcacctagatcc
ttttaaattaaaaatgaagttttaaatcaatctaaagtatatatgagtaaacttggtctgacagttaccaatgcttaatc
agtgaggcacctatctcagcgatctgtctatttcgttcatccatagttgcctgactccccgtcgtgtagataactacgat
acgggagggcttaccatctggccccagtgctgcaatgataccgcgagacccacgctcaccggctccagatttatcagcaa
taaaccagccagccggaagggccgagcgcagaagtggtcctgcaactttatccgcctccatccagtctattaattgttgc
cgggaagctagagtaagtagttcgccagttaatagtttgcgcaacgttgttgccattgctacaggcatcgtggtgtcacg
```

ctcgtcgtttggtatggcttcattcagctccggttcccaacgatcaaggcgagttacatgatcccccatgttgtgcaaaa
aagcggttagctccttcggtcctccgatcgttgtcagaagtaagttggccgcagtgttatcactcatggttatggcagca
ctgcataattctcttactgtcatgccatccgtaagatgcttttctgtgactggtgagtactcaaccaagtcattctgaga
atagtgtatgcggcgaccgagttgctcttgcccggcgtcaatacgggataataccgcgccacatagcagaactttaaaag
tgctcatcattggaaaacgttcttcggggcgaaaactctcaaggatcttaccgctgttgagatccagttcgatataaccc
actcgtgcacccaactgatcttcagcatcttttactttcaccagcgtttctgggtgagcaaaaacaggaaggcaaaatgc
cgcaaaaaagggaataagggcgacacggaaatgt

**Supplementary Table 6:** pSB1A3-convergent pTet-Tr

aggcaaaatgccgcaaaaaagggaataagggcgacacggaaatgttgaatactcatactcttcctttttcaatattattg
aagcatttatcagggttattgtctcatgagcggatacatatttgaatgtatttagaaaaataaacaaataggggttccgc
gcacatttccccgaaaagtgccacctgacgtctaagaagcgaaaaaaccccgccgaagcggggtttttttgcgcgacccgg
ctaaaaaaaaccccgccctgtcaggggcggggtttttttttgatagactccctatcagtgatagagattgacatccctat
cagtgatagagatactgagcactattggactgactattctgtgcaatagtcagtaaagcagggataaacgagatagata
agataagatagatcctaccatgattgtgctcagtatcttgttatccgctcacaatgtcaatgttatccgctcacatttat
gctagccaggcatcaaataaaacgaaaggctcagtcgaaagactgggcctttcgttttatctgttgtttgtcggtgaacg
ctctctactagagtcacactggctcaccttcgggtgggcctttctgcgtttatattacctgaagaataagtcagccgtgt
aacccgatgaggaattcttgacggctagctcagtcctaggtatagtgctagcgggtcttatcttatctatctcgtttatc
cctgcatacagaaacagaggagatacgcaatgataaacgagaacctggcggcagcgcaaaagatgagcaaaggcgaagaa
ctgttcacgggtgtggttccgatcctggttgaactggatggcgatgtgaacggtcataaatttagcgtgtctggtgaagg
cgaaggtgatgcgacctacggcaaactgacgctgaaactgatttgcaccacgggtaaactgccggttccgtggccgaccc
tggtgaccacgctgggttatggtctgatgtgtttcgcacgttacccggatcacatgaaacgccatgatttctttaaatct
gcgatgccggaaggctatgtgcaggaacgtaccatctttttcaaagatgatggtaactacaaaacccgcgcggaagttaa
atttgaaggcgatacgctggtgaaccgtattgaactgaaaggtatcgatttcaaagaagatggcaatattctgggtcaca
aactggaatacaactacaacagtcataacgtgtacattaccgccgataaacagaaaaacggtatcaaagcaaacttcaaa
atccgtcacaacatcgaagatggcggtgttcagctggccgatcattaccagcagaacaccccgattggcgatggtccggt
gctgctgccggataatcattatctgagttaccagagcaaactgtctaaagatccgaatgaaaaacgcgatcacatggttc
tgctggaatttgtgaccgcggccggcattacgcatggtatggatgaactgtataaataaccaggcatcaaataaaacgaa
aggctcagtcgaaagactgggcctttcgttttatctgttgtttgtcggtgaacgctctctactagagtcacactggctca
ccttcgggtgggcctttctgcgtttataactagtgcgctgcagtccggcaaaaaagggcaaggtgtcaccaccctgccct
ttttctttaaaaccgaaaagattacttcgcgttatgcaggcttcctcgctcactgactcgctgcgctcggtcgttcggct
gcggcgagcggtatcagctcactcaaaggcggtaatacggttatccacagaatcaggggataacgcaggaaagaacatgt
gagcaaaaggccagcaaaaggccaggaaccgtaaaaaggccgcgttgctggcgtttttccacaggctccgcccccctgac
gagcatcacaaaaatcgacgctcaagtcagaggtggcgaaacccgacaggactataaagataccaggcgtttccccctgg
aagctccctcgtgcgctctcctgttccgaccctgccgcttaccggatacctgtccgcctttctcccttcgggaagcgtgg
cgctttctcatagctcacgctgtaggtatctcagttcggtgtaggtcgttcgctccaagctgggctgtgtgcacgaaccc
cccgttcagcccgaccgctgcgccttatccggtaactatcgtcttgagtccaacccggtaagacacgacttatcgccact
ggcagcagccactggtaacaggattagcagagcgaggtatgtaggcggtgctacagagttcttgaagtggtggcctaact
acggctacactagaagaacagtatttggtatctgcgctctgctgaagccagttaccttcggaaaaagagttggtagctct
tgatccggcaaacaaaccaccgctggtagcggtggtttttttgtttgcaagcagcagattacgcgcagaaaaaaaggatc
tcaagaagatcctttgatcttttctacggggtctgacgctcagtggaacgaaaactcacgttaagggattttggtcatga
gattatcaaaaaggatcttcacctagatccttttaaattaaaaatgaagttttaaatcaatctaaagtatatatgagtaa
acttggtctgacagttaccaatgcttaatcagtgaggcacctatctcagcgatctgtctatttcgttcatccatagttgc
ctgactccccgtcgtgtagataactacgatacgggagggcttaccatctggccccagtgctgcaatgataccgcgagacc
cacgctcaccggctccagatttatcagcaataaaccagccagccggaagggccgagcgcagaagtggtcctgcaacttta
tccgcctccatccagtctattaattgttgccgggaagctagagtaagtagttcgccagttaatagtttgcgcaacgttgt
tgccattgctacaggcatcgtggtgtcacgctcgtcgtttggtatggcttcattcagctccggttcccaacgatcaaggc
gagttacatgatcccccatgttgtgcaaaaaagcggttagctccttcggtcctccgatcgttgtcagaagtaagttggcc
gcagtgttatcactcatggttatggcagcactgcataattctcttactgtcatgccatccgtaagatgcttttctgtgac
tggtgagtactcaaccaagtcattctgagaatagtgtatgcggcgaccgagttgctcttgcccggcgtcaatacgggata
ataccgcgccacatagcagaactttaaaagtgctcatcattggaaaacgttcttcggggcgaaaactctcaaggatctta
ccgctgttgagatccagttcgatataacccactcgtgcacccaactgatcttcagcatcttttactttcaccagcgtttc
tgggtgagcaaaaacagga

**Supplementary Table 7:** pSB1A3-T2AT pTet-Tr

ctcttcctttttcaatattattgaagcatttatcagggttattgtctcatgagcggatacatatttgaatgtatttagaa
aaataaacaaataggggttccgcgcacatttccccgaaaagtgccacctgacgtctaagaagacctcgcgaaaaaacccc
gccgaagcggggtttttttgcgcgaccacgacggctaaaaaaaaccccgccctgtcaggggcggggtttttttttgagcgg
gactgactattctgtgcaatagtcagtaaagcagggataaacgagatagataagataagatagatcctaccatgatttaa
acaaaattatttgtagaggactgtttcggcccttttggccatcgtcaggtcggatacacatccggcgacagtctctcga
ggactccctatcagtgatagagattgacatccctatcagtgatagagatactgagcactattgggactgactattctgtg
caatagtcagtaaagcagggataaacgagatagataagataagatagatcctaccatgattgtgctcagtatcttgttat
ccgctcacaatgtcaatgttatccgctcacatttatgctagccaggcatcaaataaaacgaaaggctcagtcgaaagact
gggcctttcgttttatctgttgtttgtcggtgaacgctctctactagagtcacactggctcaccttcgggtgggcctttc
tgcgtttatattacctgaagaataagtcagccgtgtaacccgatgaggaattcttgacggctagctcagtcctaggtata
gtgctagcgggtcttatcttatctatctcgtttatccctgcatacagaaacagaggagatacgcaatgataaacgagaac
ctggcggcagcgcaaaagatgagcaaaggcgaagaactgttcacgggtgtggttccgatcctggttgaactggatggcga

tgtgaacggtcataaatttagcgtgtctggtgaaggcgaaggtgatgcgacctacggcaaactgacgctgaaactgattt
gcaccacgggtaaactgccggttccgtggccgaccctggtgaccacgctgggttatggtctgatgtgtttcgcacgttac
ccggatcacatgaaacgccatgatttctttaaatctgcgatgccggaaggctatgtgcaggaacgtaccatctttttcaa
agatgatggtaactacaaaacccgcgcggaagttaaatttgaaggcgatacgctggtgaaccgtattgaactgaaaggta
tcgatttcaaagaagatggcaatattctgggtcacaaactggaatacaactacaacagtcataacgtgtacattaccgcc
gataaacagaaaaacggtatcaaagcaaacttcaaaatccgtcacaacatcgaagatggcggtgttcagctggccgatca
ttaccagcagaacaccccgattggcgatggtccggtgctgctgccggataatcattatctgagttaccagagcaaactgt
ctaaagatccgaatgaaaaacgcgatcacatggttctgctggaatttgtgaccgcggccggcattacgcatggtatggat
gaactgtataaataaccaggcatcaaataaaacgaaaggctcagtcgaaagactgggcctttcgttttatctgttgtttg
tcggtgaacgctctctactagagtcacactggctcaccttcgggtgggcctttctgcgtttataactagtgcgctgcagt
ccggcaaaaaagggcaaggtgtcaccaccctgccctttttctttaaaaccgaaaagattacttcgcgttatgcaggcttc
ctcgctcactgactcgctgcgctcggtcgttcggctgcggcgagcggtatcagctcactcaaaggcggtaatacggttat
ccacagaatcaggggataacgcaggaaagaacatgtgagcaaaaggccagcaaaaggccaggaaccgtaaaaaggccgcg
ttgctggcgtttttccacaggctccgcccccctgacgagcatcacaaaaatcgacgctcaagtcagaggtggcgaaaccc
gacaggactataaagataccaggcgtttccccctggaagctccctcgtgcgctctcctgttccgaccctgccgcttaccg
gatacctgtccgcctttctcccttcgggaagcgtggcgctttctcatagctcacgctgtaggtatctcagttcggtgtag
gtcgttcgctccaagctgggctgtgtgcacgaaccccccgttcagcccgaccgctgcgccttatccggtaactatcgtct
tgagtccaacccggtaagacacgacttatcgccactggcagcagccactggtaacaggattagcagagcgaggtatgtag
gcggtgctacagagttcttgaagtggtggcctaactacggctacactagaagaacagtatttggtatctgcgctctgctg
aagccagttaccttcggaaaaagagttggtagctcttgatccggcaaacaaaccaccgctggtagcggtggtttttttgt
ttgcaagcagcagattacgcgcagaaaaaaaggatctcaagaagatcctttgatcttttctacggggtctgacgctcagt
ggaacgaaaactcacgttaagggattttggtcatgagattatcaaaaaggatcttcacctagatccttttaaattaaaaa
tgaagttttaaatcaatctaaagtatatatgagtaaacttggtctgacagttaccaatgcttaatcagtgaggcacctat
ctcagcgatctgtctatttcgttcatccatagttgcctgactccccgtcgtgtagataactacgatacgggagggcttac
catctggccccagtgctgcaatgataccgcgagacccacgctcaccggctccagatttatcagcaataaaccagccagcc
ggaagggccgagcgcagaagtggtcctgcaactttatccgcctccatccagtctattaattgttgccgggaagctagagt
aagtagttcgccagttaatagtttgcgcaacgttgttgccattgctacaggcatcgtggtgtcacgctcgtcgtttggta
tggcttcattcagctccggttcccaacgatcaaggcgagttacatgatcccccatgttgtgcaaaaaagcggttagctcc
ttcggtcctccgatcgttgtcagaagtaagttggccgcagtgttatcactcatggttatggcagcactgcataattctct
tactgtcatgccatccgtaagatgcttttctgtgactggtgagtactcaaccaagtcattctgagaatagtgtatgcggc
gaccgagttgctcttgcccggcgtcaatacgggataataccgcgccacatagcagaactttaaaagtgctcatcattgga
aaacgttcttcggggcgaaaactctcaaggatcttaccgctgttgagatccagttcgatataacccactcgtgcacccaa
ctgatcttcagcatcttttactttcaccagcgtttctgggtgagcaaaaacaggaaggcaaaatgccgcaaaaaagggaa
taagggcgacacggaaatgttgaatactcata

**Supplementary Table 8:** pSB3C5-XOR (convergent pTet-Tr2 J23111-TS2-mCerulean + T2AT pLac-Tr J23111-TS1-mVenus)

tggctggtttattgctgataaatctggagccggtgagcgtgggtctcgcggtatcattgcagcactggggccagatggta
agccctcccgtatcgtagttatctacacgacggggagtcaggcaactatggatgaacgaaatagacagatcgctgagata
ggtgcctcactgattaagcattggtaactgtcagaccaagtttactcatatatactttagattgatttaaaacttcattt
ttaatttaaaaggatctaggtgaagatcctttttgataatctcatgaccaaaatcccttaacgtgagttttcgttccact
gagcgtcagaccccttaataagatgatcttcttgagatcgttttggtctgcgcgtaatctcttgctctgaaaacgaaaaa
accgccttgcagggcggtttttcgaaggttctctgagctaccaactctttgaaccgaggtaactggcttggaggagcgca
gtcaccaaaacttgtcctttcagtttagccttaaccggcgcatgacttcaagactaactcctctaaatcaattaccagtg
gctgctgccagtggtgcttttgcatgtctttccgggttggactcaagacgatagttaccggataaggcgcagcggtcgga
ctgaacggggggttcgtgcatacagtccagcttggagcgaactgcctacccggaactgagtgtcaggcgtggaatgagac
aaacgcggccataacagcggaatgacaccggtaaaccgaaaggcaggaacaggagagcgcacgagggagccgccaggga
aacgcctggtatctttatagtcctgtcgggtttcgccaccactgatttgagcgtcagatttcgtgatgcttgtcagggg
gcggagcctatggaaaaacggctttgccgcggccctctcacttccctgttaagtatcttcctggcatcttccaggaaatc
tccgccccgttcgtaagccatttccgctcgccgcagtcgaacgaccgagcgtagcgagtcagtgagcgaggaagcggaat
atatcctgtatcacatattctgctgacgcaccggtgcagcctttttttctcctgccacatgaagcacttcactgacaccct
catcagtgccaacatagtaagccagtatacactccgctagcgctgaggtctgcctcgtgaagaaggtgttgctgactcat
accaggcctgaatcgccccatcatccagccagaaagtgagggagccacggttgatgagagctttgttgtaggtggaccag
ttggtgattttgaacttttgctttgccacggaacggtctgcgttgtcgggaagatgcgtgatctgatccttcaactcagc
aaaagttcgatttattcaacaaagccacgttgtgtctcaaaatctctgatgttacattgcacaagataaaaatatatcat
catgaacaataaaactgtctgcttacataaacagtaatacaaggggtgtttactagaggttgatcgggcacgtaagaggt
tccaactttcaccataatgaaataagatcactaccgggcgtattttttgagttatcgagattttcaggagctaaggaagc
taaaatggagaaaaaaatcacgggatataccaccgttgatatatcccaatggcatcgtaaagaacattttgaggcatttc
agtcagttgctcaatgtacctataaccagaccgttcagctggatattacggcctttttaaagaccgtaaagaaaaataag
cacaagttttatccggcctttattcacattcttgcccgcctgatgaacgctcacccggagtttcgtatggccatgaaaga
cggtgagctggtgatctgggatagtgttcacccttgttacaccgttttccatgagcaaactgaaacgttttcgtccctct
ggagtgaataccacgacgatttccggcagtttctccacatatattcgcaagatgtggcgtgttacggtgaaaacctggcc
tatttccctaaagggtttattgagaatatgtttttgtctcagccaatccctgggtgagtttcaccagttttgatttaaa
cgtggccaatatggacaacttcttcgcccccgttttcacgatgggcaaatattatacgcaaggcgacaaggtgctgatgc
cgctggcgatccaggttcatcatgccgtttgtgatggcttccatgtcggccgcatgcttaatgaattacaacagtactgt
gatgagtggcagggcggggcgtaataatactagctccggcaaaaaaacgggcaaggtgtcaccaccctgccctttttctt
taaaaccgaaaagattacttcgcgtttgccacctgacgtctaagaagcgaaaaaaccccgccgaagcggggtttttttgcg
cgacccggctaaaaaaaaccccgccctgtcaggggcggggtttttttttgatagactccctatcagtgatagagattgac
atccctatcagtgatagagatactgagcactattgggacagatccactgaggcgtggatctgtgaacactaaactaaacg

```
acaatgtaatcaaactaacggatggctacttgtgtgctcagtatcttgttatccgctcacaatgtcaatgttatccgctc
acatttatgctagccaggcatcaaataaaacgaaaggctcagtcgaaagactgggcctttcgttttatctgttgtttgtc
ggtgaacgctctctactagagtcacactggctcaccttcgggtgggcctttctgcgtttatattacctgaagaataagtc
agccgtgtaacccgatgaggaattcttgacggctagctcagtcctaggtatagtgctagcgggagtttgattacattgtc
gtttagtttagtgatacataaacagaggagatatcacatgactaaacgaaacctggcggcagcgcaaaagatggtgagta
aaggcgaagagctgttcacaggggttgttccgattctggtcgaactggacggggacgttaatggtcacaaattcagcgtt
agcggtgagggcgagggtgatgccacttatggtaaactgaccctgaaattcatctgtaccaccggcaaactgcctgttcc
ttggcctacactggttacaacactgacttggggtgttcaatgttttgctcgctatccggatcacatgaaacagcacgatt
tcttcaaaagcgccatgcctgaaggttatgtccaagagcgtacgatcttctttaaagacgacggcaactataaaacccgt
gccgaggtgaaattcgaaggtgataccctggtaaaccgtatcgaactgaaagggatcgacttcaaagaggacgggaacat
tctgggccataaactggagtataacgccatcagcgataatgtgtatattaccgccgacaaacagaaaaacgggatcaaag
ccaacttcaaaatccgccacaacatcgaggatggtagcgttcaactggccgatcactatcaacagaataccccgattggt
gatggtcctgttctgctgcctgataaccactatctgagcacccagtctaaactgtccaaagacccgaacgagaaacgtga
tcacatggttctgctggagtttgttaccgctgccggcattactctgggtatggatgaactgtataaatagcataacccct
tggggcctctaaacgggtcttgaggggttttttgctgcagtccggcaaaaaagggcaaggtgtcaccaccctgcccttt t
tctttaaaaccgaaaagattacttcgcgttatgcaggcttcctcgctcactgactcgctgcgctcggtcgttcggctgcg
gcgagcggtatcagctcactaagcttgacctcgcgaaaaaccccgccgaagcggggtttttgcgcgaccacgacggct
aaaaaaaccccgccctgtcaggggcggggtttttttttgagcggactgactattctgtgcaatagtcagtaaagcagg
gataaacgagatagataagataagatagatcctaccatgatttaaacaaaattatttgtagaggactgtttcggcccttt
tgggccatcgtcaggtcggatacacatccggcgacagtctctcgagataaatgtgagcggataacattgacattgtgagc
ggataacaagatactgagcacagggactgactattctgtgcaatagtcagtaaagcagggataaacgagatagataagat
aagatagatcctaccatgatgtgctcagtatctctatcactgatagggatgtcaatctctatcactgatagggagagacc
aggcatcaaataaaacgaaaggctcagtcgaaagactgggcctttcgttttatctgttgtttgtcggtgaacgctctcta
ctagagtcacactggctcaccttcgggtgggcctttctgcgtttatagtgcgctgacaagtccattaccaaggagtgcga
tgcagttactcagaattcttgacggctagctcagtcctaggtatagtgctagcgggtcttatcttatctatctcgtttat
ccctgcatacagaaacagaggagatacgcaatgataaacgagaacctggcggcagcgcaaaagatgagcaaaggcgaaga
actgttcacgggtgtggttccgatcctggttgaactggatggcgatgtgaacggtcataaatttagcgtgtctggtgaag
gcgaaggtgatgcgacctacggcaaactgacgctgaaactgatttgcaccacgggtaaactgccggttccgtggccgacc
ctggtgaccacgctgggttatggtctgatgtgtttcgcacgttacccggatcacatgaaacgccatgatttctttaaatc
tgcgatgccggaaggctatgtgcaggaacgtaccatctttttcaaagatgatggtaactacaaaacccgcgcggaagtta
aatttgaaggcgatacgctggtgaaccgtattgaactgaaaggtatcgatttcaaagaagatggcaatattctgggtcac
aaactggaatacaactacaacagtcataacgtgtacattaccgccgataaacagaaaaacggtatcaaagcaaacttcaa
aatccgtcacaacatcgaagatggcggtgttcagctggccgatcattaccagcagaacaccccgattggcgatggtccgg
tgctgctgccggataatcattatctgagttaccagagcaaactgtctaaagatccgaatgaaaaacgcgatcacatggtt
ctgctggaatttgtgaccgcggccggcattacgcatggtatggatgaactgtataaataaccaggcatcaaataaaacga
aaggctcagtcgaaagactgggcctttcgttttatctgttgtttgtcggtgaacgctctctactagagtcacactggctc
accttcgggtgggcctttctgcgtttataactagtagcggccgctgcaggagtcactaagggttagttagttagattagc
agaaagtcaaaagcctccgaccggaggcttttgactaaaacttcccttggggttatcattggggctcactcaaaggcggt
aatcagataaaaaaaatccttagctttcgctaaggatgatttctgctagagatggaatagactggatggaggcggataaa
gttgcaggaccacttctgcgctcggcccttccggc
```


1. Lutz, R.; Bujard, H., Independent and tight regulation of transcriptional units in Escherichia coli via the LacR/O, the TetR/O and AraC/I1-I2 regulatory elements. *Nucleic Acids Res* **1997,** *25* (6), 1203-10.

2. http://parts.igem.org/Part:BBa_K783076; Group: iGEM12_BostonU (2012-10-03)
